# Early-life stress and ovarian hormones alter transcriptional regulation in the nucleus accumbens resulting in sex-specific responses to cocaine

**DOI:** 10.1101/2023.04.14.536984

**Authors:** Devin Rocks, Ivana Jaric, Fabio Bellia, Heining Cham, John M. Greally, Masako Suzuki, Marija Kundakovic

## Abstract

Early-life stress and ovarian hormones contribute to increased female vulnerability to cocaine addiction. Here we reveal molecular substrates in the key reward area, the nucleus accumbens, through which these female-specific factors affect immediate and conditioning responses to cocaine in mice. We find shared involvement of X chromosome and estrogen signaling gene regulation in enhanced conditioning responses seen after early-life stress and during the low-estrogenic state in females. During the low-estrogenic state, females respond to acute cocaine exposure by increasing the accessibility of neuronal chromatin enriched for the binding sites of ΔFosB, a transcription factor implicated in chronic cocaine response and addiction. Conversely, high-estrogenic females respond to cocaine by preferential closing of neuronal chromatin, providing a mechanism for limiting cocaine-driven chromatin and synaptic plasticity. We find that physiological estrogen withdrawal, exposure to early-life stress, and absence of the second X chromosome all nullify the protective effect of high-estrogenic state on cocaine conditioning in females. Our findings offer a molecular framework to understand sex-specific neuronal mechanisms underlying cocaine use disorder.

## Introduction

Substance use disorders affect people across genders, although a significant body of evidence shows that women and men develop these disorders differently. Specifically, cocaine use disorder (CUD) is a major global health issue which affects ∼1 million individuals over age 12 in the United States alone, (1) and there is overwhelming evidence that females are more sensitive to the effects of this drug (2). Women are reported to transition to addiction faster, have more difficulty remaining abstinent, and experience more adverse consequences of cocaine use (3). While there are studies showing that the ovarian hormone estradiol potentiates the cocaine-induced “high” and makes women more sensitive to cocaine (4, 5), very few studies have addressed the underlying molecular mechanism. In fact, animal studies have been traditionally focused on males (6), missing on the opportunity to reveal sex-specific mechanisms underlying reward processing and cocaine addiction.

Clinical studies including childhood trauma survivors offer some insights into the increased female vulnerability to cocaine. While early-life trauma is a major risk factor for the development of mental disorders in both males and females (7), the association between childhood trauma and the development of substance use disorders is generally stronger in women. In fact, childhood traum was shown to increase the likelihood of cocaine relapse and drug use escalation in women but not in men (8). Revealing sex-specific and trauma-related pathophysiology of cocaine addiction can be of great benefit in improving CUD treatment outcomes in people of all genders.

In terms of molecular mechanisms, changes in chromatin and gene expression in the key brain reward area, the nucleus accumbens (NAc), have long been proposed to underlie both the acute and chronic effects of cocaine (9). However, the sex specificity of cocaine’s effects is understudied. We recently showed that neuronal chromatin organization, a major mechanism controlling gene expression, changes in the ventral hippocampus with sex and the estrous cycle stage (10). Earlier studies have also shown transcriptional and chromatin changes in the NAc in response to early-life stress (11–13). Therefore, we hypothesized that early-life stress and ovarian hormone status can affect transcriptional and chromatin regulation in the NAc leading to sex specific responses to cocaine.

Here, we designed a comprehensive mouse study to address sex difference in cocaine sensitivity at both the etiological and mechanistic level. We first demonstrate the effects of the two major sex-specific risk factors in the etiology of CUD, early-life stress and ovarian hormone status, on the acquisition of cocaine preference in mice. We then characterize the NAc transcriptome and epigenome to reveal both shared and distinct transcriptional mechanisms by which these two factors drive sex-specific responses to cocaine, including an unexpected involvement of the inactive X chromosome which we functionally linked to cocaine-induced behavior.

## Results

### Early life stress study design

To determine the effects of early-life stress on addiction-related behavioral and molecular phenotypes, we used a previously established maternal separation paradigm (14–16). Mouse litters were assigned to either the control (Con) group that received no stress or an early-life stress group that underwent a 3-hour maternal separation (MS) daily from postnatal day (P) 1-14 (**Figure 1A**). During separation from their pups, dams were also subjected to unpredictable stress (forced swim or restrain stress) to prevent the occurrence of compensatory maternal care following separation (17). We monitored maternal behaviors during the first six days postpartum (P1-6), known to be the critical period for mother-infant interactions in mice (18), and confirmed that there was a disruption in maternal care for at least one hour following separation (**Figure 1B**). Specifically, we found a reduction in total nurturing behaviors including nursing, arched-back nursing, and licking and grooming in MS compared to control mothers across the six days (F(_1, 270_)_=_ 12.44, p=0.01; Repeated-measures ANOVA) and on average (t(_45_)=7.15, p=5.9e-09; Welch Two Sample t-test). We also found an increase in the time spent out of the nest in MS mothers compared to control mothers across the six days (F(_1,270_)=10.18, p=0.002; Repeated-measures ANOVA) and on average (t(_45_)=8.58, p=4.4e-11; Welch Two Sample t-test). Animals were weaned at P28 and the majority of Con and MS female and male animals underwent behavioral testing using the cocaine-induced conditioned place preference (CPP) test during the late adolescent period in mice, from P54-P60 (**Figure 1A**). A subset of Con and MS male and female mice were sacrificed at P50, just prior to the onset of CPP, and used for RNA-seq analysis (**Figure 1A**), to assess MS-induced gene expression changes in the NAc that may underlie the early-life stress-induced CPP phenotype.

**Figure 1.**
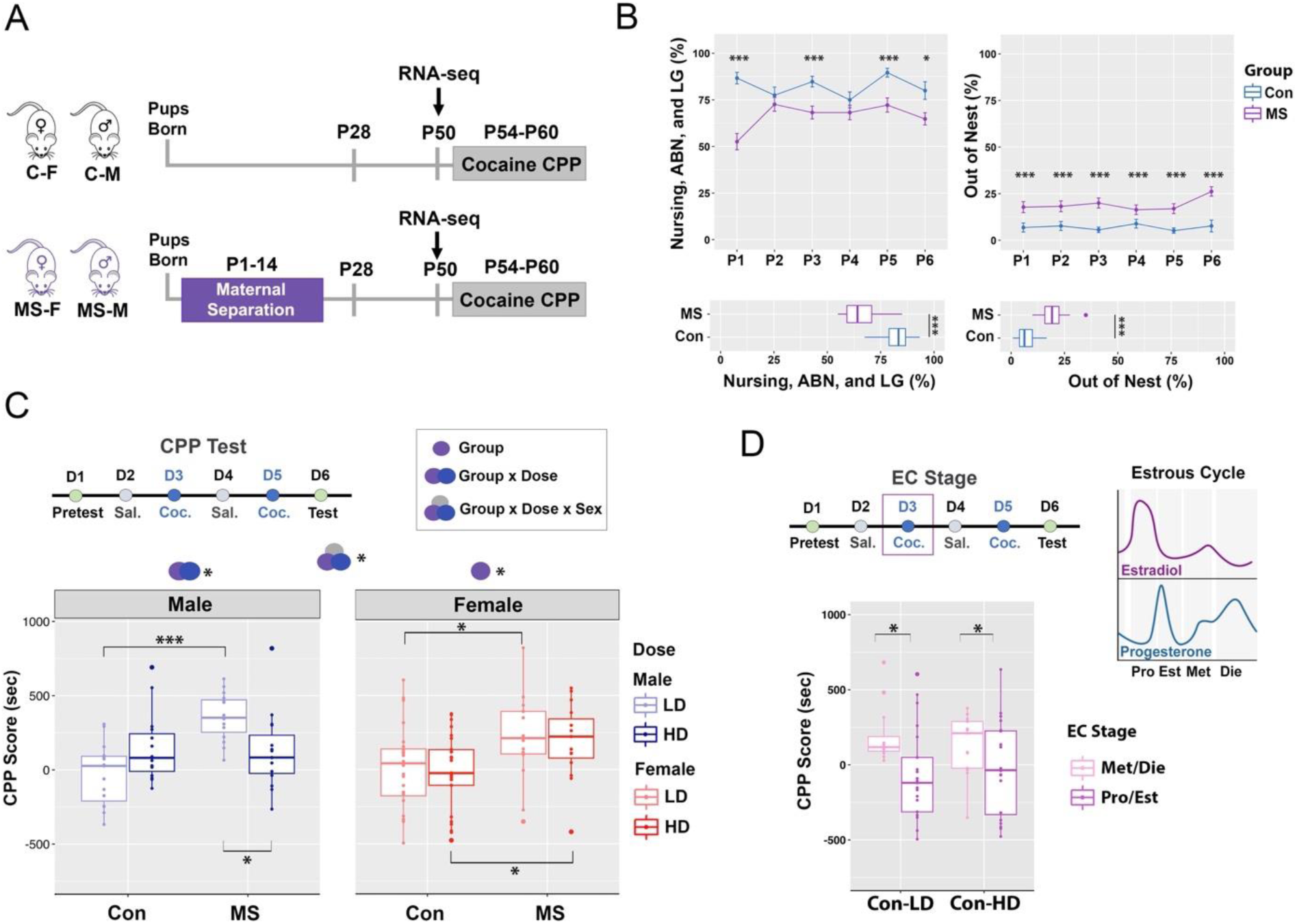
Early-life stress alters cocaine preference in a sex-and dose-dependent manner. **(A)** The early-life stress study included two major experimental groups:control (Con) mice and mice that underwent a maternal separation (MS) paradigm from postnatal day (P) 1-14. Female (F) and male (M) mouse offspring were weaned at P28 and either underwent cocaine CPP from P54-P60 or were euthanized for RNA-seq analysis at P50. **(B)** Maternal behavior was scored 1h following maternal separation, including the time spent nursing, arched-back nursing (ABN), and licking and grooming (LG) across P1-6 (upper-left) and on average (lower-left), as well the time spent out of the nest across P1-P6 (upper-right) and on average (lower-right). *p < 0.05; **p < 0.01; ***p < 0.001; Repeated-measures ANOVA with the Tukey’s post hoc test (upper plots); Welch Two-sample T-test (lower plots). **(C)** The cocaine place preference (CPP) test spanned 6 days (D) and cocaine was administered on D3 and D5, at either low dose (LD; 2.5 mg/kg) or high dose (HD; 10 mg/kg). CPP scores were analyzed in all animals with group, sex, and dose as factors; (n = 15-25 animals/sex/dose/group). Symbols above the graphs show significant main effects of group, sex, dose, or their interaction. **(D)** The CPP score analysis was repeated in control female mice considering their estrous cycle stage (depicted above) on the first day of cocaine exposure. This analysis was done with dose and estrogen status as factors; (n = 12–23/stage/dose). *p < 0.05; **p < 0.01; ***p < 0.001; Three-way ANOVA (Group x Dose x Sex interaction); Two-way ANOVA (male, female, and estrous cycle plots). Box plots (box, 1st–3rd quartile; horizontal line, median; whiskers, 1.5× IQR). C-F, control female; C-M, control male; MS-F, maternal separation female; MS-M, maternal separation male; Sal., saline; Coc., cocaine; Pro, proestrus; Est, estrus; Met, metestrus; Die, diestrus.

### Early-life stress alters cocaine preference in a sex-and dose-dependent manner

To evaluate the effects of early-life stress on cocaine preference, we performed the CPP test using two different cocaine doses:the higher dose (HD, 10 mg/kg, i.p.) shown to effectively induce cocaine preference (19) and the lower dose (LD, 2.5 mg/kg, i.p.) considered to be insufficient to induce CPP under basal conditions (19, 20). HD or LD cocaine was administered to male and female mice of Con and MS groups on conditioning days (Days 3 and 5, **Figure 1C**). To account for the effect of the estrous cycle stage on CPP score in females, we performed estrous cycle tracking daily across the entire experiment (Days 1-6).

We first analyzed the CPP score using a three-way ANOVA with experimental group (Con or MS), sex (female or male), and dose (HD or LD) as factors (**Figure 1C**, **Supplementary Table 1A**). Importantly, we observed a significant group by sex by dose interaction effect (F_(1, 132)_= 4.32, p=0.04; three-way ANOVA). To further understand this three-way effect, we analyzed the effect of group and dose, and their interaction, separately in each sex (**Supplementary Table 1A)**. Within males, we found that the effect of early-life stress depends on dose (group x dose interaction; F_(1, 132)_= 7.51, p=0.01; two-way ANOVA; **Figure 1C**). The high cocaine dose induced a preference for the cocaine-paired compartment in both control (Con-HD) and early-life stress (MS-HD) male groups, with no significant difference between these two groups (F_(1, 132)_= 0.03, p=0.85, **Figure 1C**). However, we found a significant effect of early-life stress with the lower dose of cocaine (F_(1, 132)_= 13.62, p=0.0003; **Suppl. Table 1A**), where males that underwent early-life stress (MS-LD) showed cocaine preference that was not observed in the control (Con-LD) male group. Interestingly, within MS males, the LD group had a higher CPP score than the HD group (F(_1, 132_)= 4.89, p=0.03), indicating that early-life stress reversed the expected dose-response relationship in male animals (**Figure 1C**).

Within females, we observed a significant effect of group (F_(1, 132)_= 4.78, p=0.03; two-way ANOVA), with no effect of dose and no group by dose interaction (**Figure 1C, Suppl. Table 1A**). Further analysis within females revealed a significant main effect of group in both the LD (F_(1, 132)_= 4.46, p=0.04) and HD (F_(1, 132)_= 4.78, p=0.03) groups, with females exposed to early-life stress exhibiting a higher cocaine-induced preference than control females regardless of cocaine dose.

Notably, unlike males, control females receiving the high cocaine dose (Con-HD), showed no overall preference for the cocaine-paired compartment, with an average CPP score of-17.50 (*±* = 256.23), and responses of individual animals that varied from preference to aversion (**Figure 1C**). Based on the known effect of estrogen on cocaine-induced dopamine levels in the brain reward area (2), we hypothesized that the varied CPP score in control female animals may be explained by their sex hormone status. An important point of consideration, though, is that the CPP test lasts six days and the hormone status at any of the days (Days 1-6) could influence animal performance on the day of test (Day 6). We noticed that cocaine may affect estrous cycling, consistent with previous literature (21) so, for our analysis, we decided to focus on the animal’s estrous cycle stage on their first exposure to cocaine (Day 3, **Figure 1D**). To increase the power of the analysis, females were classified into two groups:1) Pro/Est animals (occupying the proestrus, estrus, or transitionary phases) with higher levels of estrogen; and 2) Met/Die animals (occupying the metestrus, diestrus, or transitionary phases) with lower levels of estrogen. We analyzed the control female CPP data using two-way ANOVA with the estrous cycle stage (Pro/Est or Met/Die) and dose (HD or LD) as factors (**Figure 1D**, **Suppl. Table 1B**). We found no effect of the estrous cycle, dose, or their interaction within control females (**Suppl. Table 1B**). After dropping the interaction effect from the model, however, we found a significant effect of the estrous cycle stage (F(_1, 64_)= 4.31, p=0.04; ANOVA), with the Met/Die control group exhibiting higher CPP scores than the Pro/Est control group across the two doses (**Figure 1D**). These data indicated that high estrogen levels in the Pro/Est group may be a protective factor against the conditioning effects of cocaine (**Figure 1D**) and this effect seems to be lost following early-life stress (**Figure 1C**).

Overall, our findings indicate that early-life stress in the form of maternal separation over the first two weeks of life increases cocaine preference in late adolescence in both male and female mice. However, the effect of early-life stress is stronger in females, increasing cocaine preference across the two doses, while having an effect with only the low cocaine dose in males.

### Transcription in the NAc is altered by early-life stress in a sex-specific manner

To determine the transcriptional signatures of early-life stress that preceded the observed cocaine-induced behavioral phenotypes, we performed RNA-seq on the NAc of MS and Con males and females at P50 (N = 6/group/sex; **Figure 2A**). To account for the effect of the estrous cycle stage on gene expression in female animals (15), we ensured that both Con and MS female groups have a balanced number of high and low estrogenic animals (see *Methods*). We first looked at the control males and females and found 1508 genes (P_adj_ < 0.10) that show sex-specific expression in the NAc under baseline conditions, including genes relevant for the regulation of chromatin and transcription, response to stress, and synaptic function (**Suppl. Figure 1**). Importantly, the number of sex-specific genes decreased to 163 following early-life stress (P_adj_ < 0.10; **Suppl. Figure 1**), implying that early-life stress reduces sex differences in the NAc, consistent with the CPP phenotype that becomes more similar between males and females after exposure to early-life stress **(Figure 1C, Suppl. Table 1)**.

**Figure 2.**
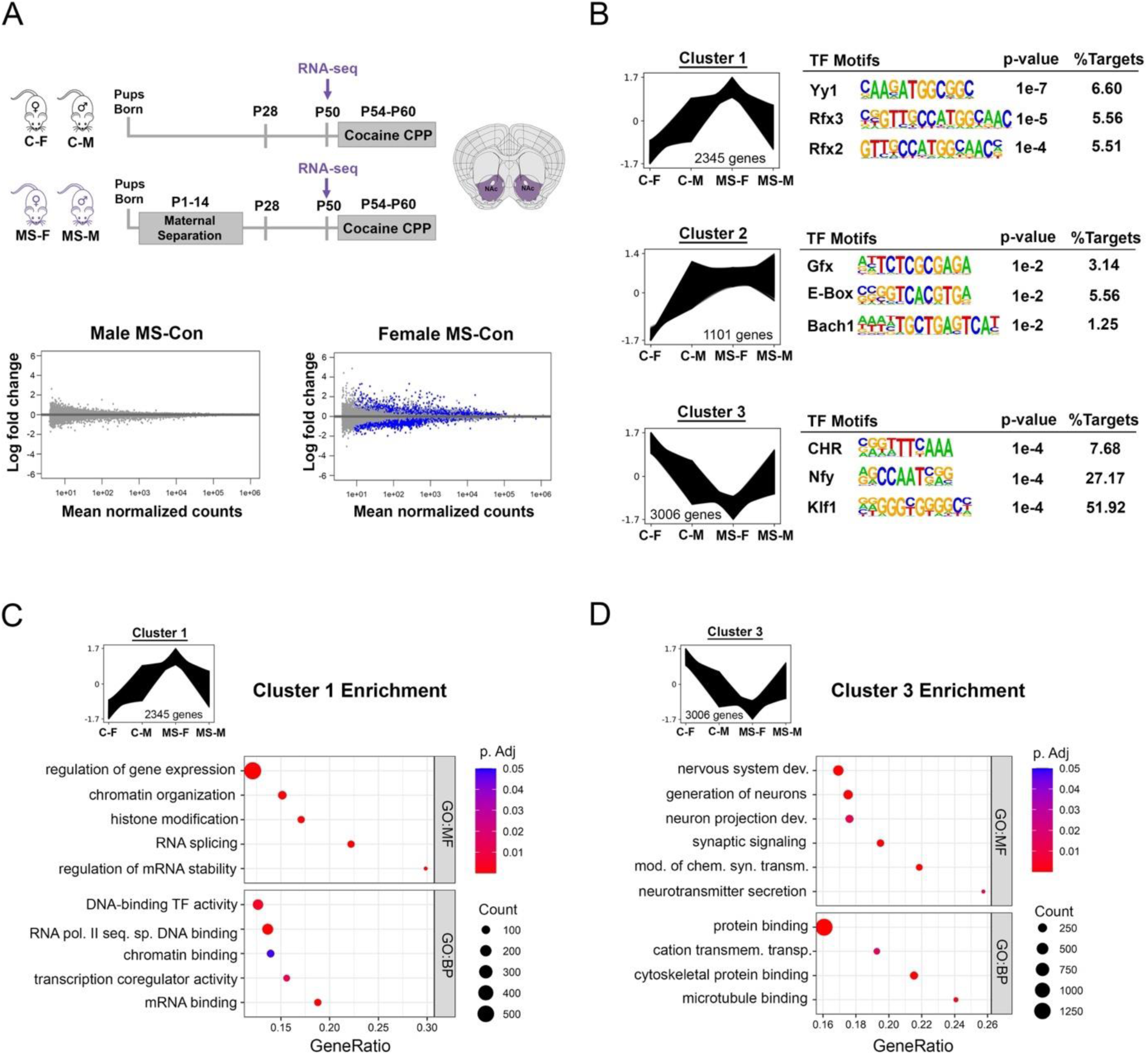
Early-life stress results in sex-specific changes in NAc gene expression. **(A)** RNA-seq was performed on the NAc from female and male mice of the control (Con) and maternal separation (MS) group (n = 6 animals/sex/group). Volcano plots depict differentially expressed genes across groups within males (n = 0) and females (n = 5527); blue dots represent significant genes (p_adj_ < 0.1). **(B)** The three significant gene clusters identified using Clust across the four experimental conditions (Con females, Con males, MS females, and MS males) are shown (left) along with the top 3 enriched transcription factor motifs in the promoter regions of cluster genes (right). **(C-D)** Dotplots depict select gene ontology (GO) terms for biological process (BP) and molecular function (MF) significantly enriched for cluster 1 (**C**) and cluster 3 (**D**) genes. Colors indicate adjusted *p-*values and dot size corresponds to gene count; Box plots (box, 1st–3rd quartile; horizontal line, median; whiskers, 1.5× IQR). C-F, control female; C-M, control male; MS-F, maternal separation female; MS-M, maternal separation male.

We then performed differential gene expression analysis between MS and Con in each sex, and we separately tested for group by sex interaction (**Figure 2A**, **Suppl. Figure 2**). Remarkably, differential gene expression analysis between MS and Con revealed 5527 differentially expressed genes (DEGs) in females, while no DEGs were identified in males using a P_adj_ < 0.10 cutoff. 455 DEGs were identified as having a significant group by sex interaction; these genes were enriched for metabolic and brain disease-relevant terms (**Suppl. Figure 2**). We compared our dataset to that of two other studies that performed RNA-seq on the NAc of male and female mice exposed to early-life stress (11, 12) (**Suppl. Figure 3**). Although our three studies used different early-life stress paradigms, the common finding was that early stress affects female gene expression more profoundly, with our MS paradigm showing the most extreme sex difference. Specifically, when we used the looser cutoff criteria that corresponded to the other two studies (11, 12), we found 15 female DEGs that overlapped between the three paradigms, showing enrichment of genes relevant to neuronal development and neurogenesis (**Suppl. Figure 3A-B**). With the same criterion, in males, we identified a similar number of DEGs as the other two studies, but there were no overlapping genes between the three datasets (**Suppl. Figure 3A**), indicating subtler and less consistent changes in male NAc gene expression following early-life stress. Interestingly, in the overlap between our female data with the female data derived from the ELS paradigm that included the combined maternal separation and limited bedding from P10-17 (11), we found genes enriched for addiction-relevant pathways (**Suppl. Figure 3C**). With the limited bedding and nesting (P2-10) paradigm (12), early-life stress altered glutamatergic-, transcription-and reproductive axis-relevant genes that overlapped with our data (**Suppl. Figure 3D**).

To include both females and males in our analysis of the RNA-seq data and address sex differences, we leveraged analysis methods that allowed us to identify broad patterns of gene expression differences across the four experimental groups. We first performed gene coexpression clustering analysis using Clust (22), which takes into account the expression of all detected transcripts and allows us to identify genes with similar changes in expression patterns across groups. We identified three distinct clusters of genes with covarying expression across the four conditions (**Figure 2B, Suppl. Table 2F**). *Cluster 1* contains 2345 genes that, on average, are more highly expressed in male than female controls. This sex difference is reversed in the MS group due to an upregulation of expression in females and lack of a transcriptional response in males. Motif analysis on this cluster revealed the top motif as the binding site for Yy1 (**Figure 2B**), a transcription factor previously implicated in mediating the transcriptional effects of the stress response (23). Though Yy1 is predicted to regulate the cluster 1 genes that are upregulated in MS females, Yy1 expression itself is downregulated in this group (**Suppl. Figure 4A, Suppl. Table 2A**), possibly as a result of its transcriptional autoregulation (24). *Cluster 2* contains 1101 genes with the top Gfx motif; these genes exhibit a similar expression pattern to cluster 1, except for a relatively reduced upregulation in MS females, making females more similar to males (**Figure 2B**). *Cluster 3*, by contrast, contains 3006 genes which are more highly expressed in females than males in the control group. This sex difference is also reversed after early-life stress due to downregulation in females. Motif analysis revealed that more than half of the genes in this cluster harbor binding sites for the Klf family of transcription factors proximal to their promoter (**Figure 2B**). Several Klf transcription factors are downregulated in MS females (**Suppl. Figure 4B, Suppl. Table 2A**), including Klf9, which has been implicated in blunting neuronal plasticity during stress exposure (25). To understand the distribution of these genes across different NAc cell types, we used a single-cell RNA-seq dataset from the rodent NAc (26). We found a biased expression across the cell types, with genes in Cluster 1 showing enriched expression in Drd1-and Drd2-medium spiny neurons (MSNs), genes in Cluster 2 showing enriched expression in astrocytes and interneurons, and genes in Cluster 3 showing enriched expression in glutamatergic neurons and interneurons (**Suppl. Figure 5**).

Given clusters 1 and 3 contain the highest number of genes, represent the highest degree of expression change, and are putatively regulated by stress-related transcription factors whose expression are altered in MS females, we focused our following analysis on these clusters. When we performed Gene Ontology (GO) enrichment analysis on cluster 1, which represents genes upregulated in MS females, we found that these genes are broadly involved in gene regulation, chromatin organization and RNA processing (**Figure 2C**), indicating that early-life stress leads to upregulation of genes related to transcriptional control in the female NAc. In cluster 3, which represents genes downregulated in MS females, GO enrichment analysis revealed terms related to neurotransmission and synaptic function, indicating a downregulation of genes important for neuronal function in the female NAc following early-life stress (**Figure 2D**).

### Early-life stress alters gene expression related to estrogen signaling and dosage compensation in females

To further identify the biological pathways affected by sex and early-life stress in the NAc, we performed gene set enrichment analysis (GSEA, (27)) on the ranked gene lists from three comparisons:MS vs. Con males, MS vs. Con females, and a group by sex interaction analysis (**Figure 3A**). By utilizing a ranked list of all detected genes, rather than relying on the smaller group of significant genes, this analysis allowed us to incorporate gene expression data from males and to continue focusing on sex differences and expression patterns across the experimental conditions. GSEA revealed pathways affected by early-life stress in males, including signaling by Ntrk2 and regulation of short-term neuronal plasticity (**Figure 3A, Suppl. Table 3**). As expected, more pathways were identified in females and in the group by sex interaction analysis than in males, including pathways related to synaptic vesicle regulation, neurotransmission, and axonogenesis (**Figure 3A, Suppl. Table 3**). Among female-specific pathways, we found that expression of genes belonging to the nuclear estrogen receptor beta (ERβ) network was affected by early-life stress. Given the role of ERβ in mediating the rewarding properties of cocaine in female mice (4), we further investigated which genes in this network were perturbed by early-life stress in female mice. We found that 5/13 genes in the ERβ network exhibited altered expression after MS (**Figure 3B**), and, more specifically, all five genes are downregulated in MS females (**Figure 3C, Suppl. Table 2A**). A deficit in estrogen receptor signaling in the NAc of MS females is consistent with their increased cocaine preference, given the protective effect of high estrogen on cocaine CPP we observed in control females (**Figure 1D**).

**Figure 3.**
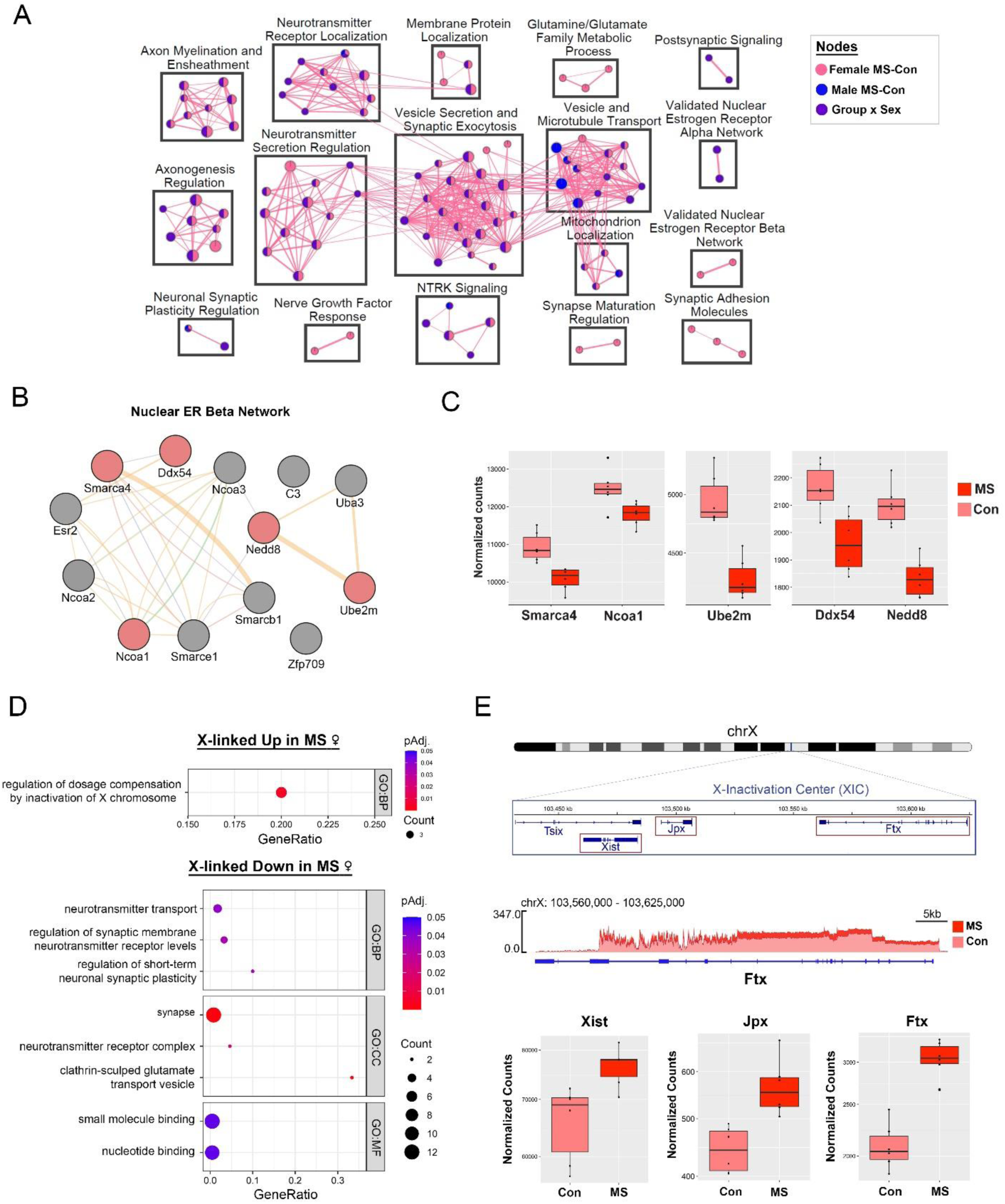
Early-life stress affects NAc gene expression related to estrogen signaling and X chromosome regulation in females. **(A)** Gene-set enrichment analysis (GSEA) was performed on ranked gene lists from the RNA-seq data in maternal separation (MS) vs. control (Con) conditions in females, males, and the group x sex interaction. Each node in the enrichment map represents a pathway significantly enriched in one or more gene lists (denoted by node color), while edges represent genes shared between pathways. Pathway clusters and their names were assigned by the AutoAnnotate app in Cytoscape. **(B)** The Nuclear Estrogen Receptor (ER) Beta Network pathway, which was significantly enriched in the female MS vs. Con gene list, was visualized using GeneMania, with each circle representing a gene in the pathway and leading-edge genes colored in green. Edges represent protein-protein interactions between pathway gene products. **(C)** Normalized count plots are shown for each of the leading-edge genes depicted in **(B)** for the female MS vs. Con comparison. **(D)** Dotplots show the only gene ontology (GO) term enriched in X-linked genes upregulated in MS females (top) and selected GO terms for biological process (BP), molecular function (MF), and cellular component (CC) enriched in X-linked genes downregulated in MS females (bottom). Colors indicate adjusted *p-*values and dot size corresponds to gene count. **(E)** A schematic of the X chromosome is shown with an inset corresponding to a portion of the X-Inactivation Center (XIC) which contains the long non-coding RNAs *Xist*, *Jpx*, and *Ftx* (top). Expression of *Ftx* between the female Con and MS groups is shown with a SparK plot of group-average normalized RNA-seq reads across the *Ftx* locus (middle, n = 6 replicates/group). Normalized count plots show expression of *Xist*, *Jpx*, and *Ftx* in the female MS vs. Con comparison (bottom). Box plots (box, 1st–3rd quartile; horizontal line, median; whiskers, 1.5× IQR).

Change in the expression of genes related to sex hormone signaling can certainly provide a molecular basis for the sex differences in behavior we observed. However, another possible mechanism involves differences in the expression of sex chromosome-linked genes between males and females. We therefore performed GO enrichment analysis on X-linked genes whose expression is altered (at p_adj_ < 0.1) by early-life stress in females (**Figure 3D**). Intriguingly, X-linked genes upregulated in MS females were enriched for only one term:regulation of dosage compensation by inactivation of X chromosome (**Figure 3D**). By contrast, several terms were identified in X-linked genes downregulated by early-life stress, including terms related to synapse, glutamate signaling, and synaptic plasticity (**Figure 3D**). These results mirror those that we identified in our cluster analysis, namely the upregulation of genes involved in transcriptional control and downregulation of genes involved in neuronal function in MS females. Remarkably, among the X-linked genes upregulated in MS females were three long non-coding RNAs (lncRNAs) that are present in the X-inactivation center (XIC, (28)) and play integral roles in the initiation and maintenance of X-inactivation:*Xist* (29), *Jpx* (30), and *Ftx* (31) (**Figure 3E**). In addition to these XIC lncRNAs, early-life stress altered the expression of several other genes, both X-linked and autosomal, involved with various aspects of X-chromosome inactivation (**Suppl. Figure 6A**) including *Xist* m^6^a methylation (*Rbm15*, *Rbm15b*, and *Ythdc1*, (32)), inactive X (Xi) localization (*Firre*, (33)), and Xi chromatin repression (*Smchd1* (34) and *Lrif1* (35)). Downregulated X-linked genes in MS females include those involved in the glutamate receptor complex (*Porcn* (36), *Dlg3* (37, 38), and *Iqsec2* (39)) and synaptic vesicle regulation (*Cdk16* (40), *Syn1* (41), *Syp* (42), and *Gdi1* (43), **Suppl. Figure 6B**). While we are unable to resolve the relative contributions of the active and inactive X chromosomes to the observed changes in gene expression, it is noteworthy that several genes downregulated in MS females are those that have been observed to escape from X-inactivation in the mouse brain (44), including *Eif2s3x*, *Gdi1*, and *Syp* (**Suppl. Table 2A**).

Taken together, early-life stress impacts the expression of genes involved in estrogen signaling and X-linked dosage compensation in the female NAc, both of which are likely to contribute to the sex-specific CPP phenotype that we observed. Considering the significant X chromosome dynamics in response to sex hormone changes across the estrous cycle (45), it is plausible that the two female-specific factors, ovarian hormone status and X chromosome regulation, may interact to shape sex-specific brain regulation under basal conditions as well as in response to environmental factors such as stress and cocaine exposure.

### Acute cocaine exposure leads to neuronal chromatin reorganization in the NAc

While the impact of ovarian hormones on cocaine’s reinforcing effect in the NAc has been known for a long time (5), the underlying molecular mechanisms have not been explored. Our CPP data indicated that the estrous cycle state at the first cocaine exposure affects the acquisition of the preference for the cocaine-paired compartment, making the female response to cocaine more variable compared to that of males at 10 mg/kg cocaine dose under basal conditions (**Figure 1C-D**). Therefore, we explored a transcriptional mechanism in NAc neurons through which acute cocaine treatment induces the observed estrous cycle-and sex-specific effects.

Based on the previous results in the ventral hippocampus showing that neuronal chromatin organization varies with the estrous cycle stage and sex (10), we hypothesized that acute cocaine exposure would differentially affect neuronal chromatin in the NAc of males and females across different phases of the cycle. To test this hypothesis, we performed ATAC-seq on NAc neuronal nuclei, purified with fluorescence-activated nuclear sorting (FANS), from proestrus (high-estrogen, low-progesterone) females, diestrus (low-estrogen, high-progesterone) females, and male mice one hour after exposure to 10 mg/kg cocaine alongside group-matched controls (**Figure 4A**).

**Figure 4.**
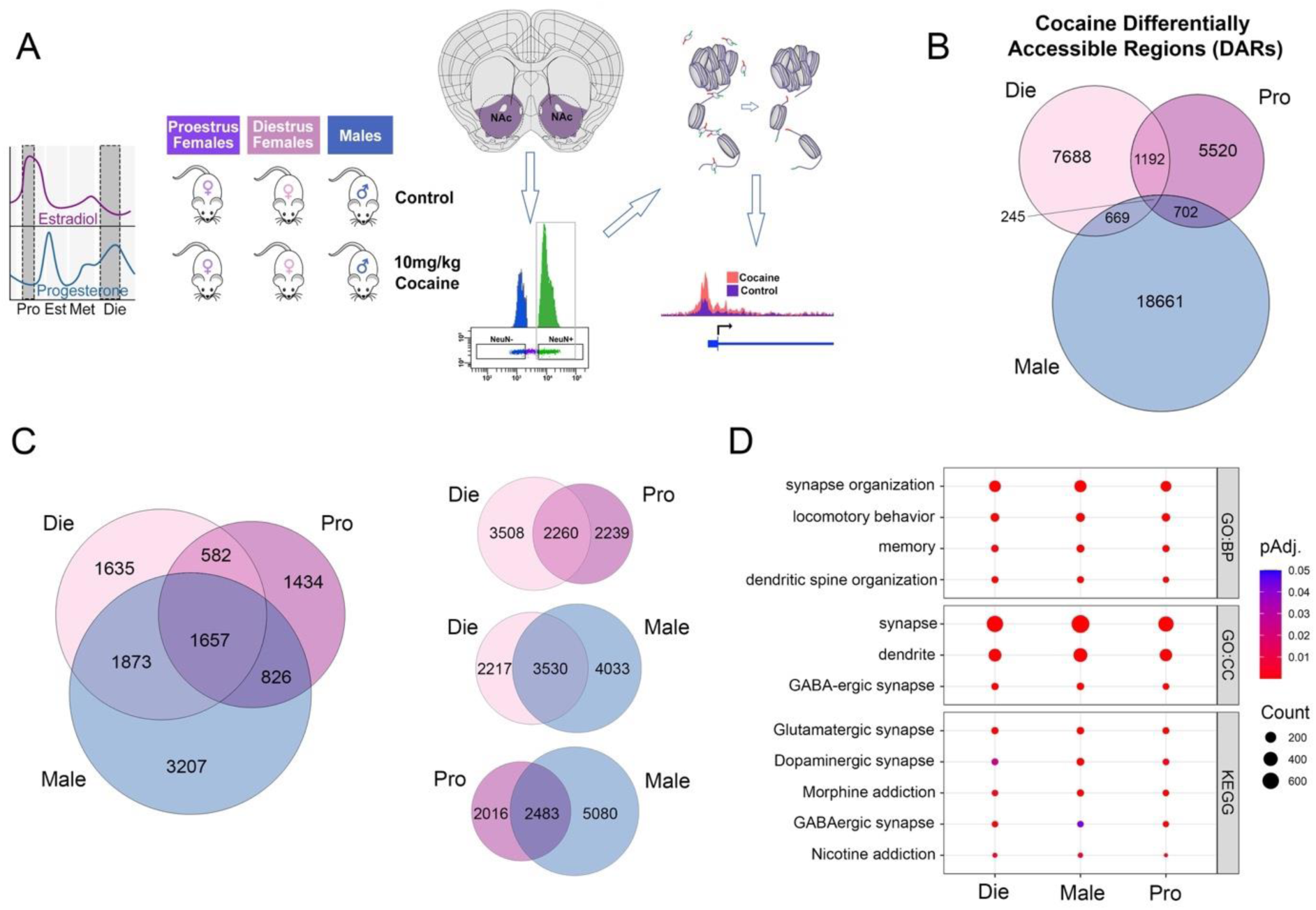
Acute cocaine exposure alters chromatin accessibility in NAc neurons. **(A)** The ATAC-seq assay was performed on FACS-purified neuronal (NeuN+) nuclei derived from the NAc of proestrus females, diestrus females, and males under basal conditions (Control) and after acute cocaine treatment (1 hour; 10 mg/kg dose). Proestrus and diestrus phases of the estrous cycle are highlighted with the corresponding estradiol and progesterone levels. **(B)** A Venn diagram shows the overlap of cocaine-induced differentially accessible regions (DARs) among the three groups. **(C)** Also shown is the overlap of genes annotated to cocaine-induced DARs across the three groups (left) and in each group comparison (right). **(D)** For each group, dot plots show select gene ontology (GO) terms for biological process (BP) and cellular component (CC), as well as KEGG pathways enriched in genes annotated to cocaine-induced DARs. Colors indicate adjusted *p-*values and dot size corresponds to gene count. Die (light pink), diestrus; Pro (purple), proestrus; Male (blue), males.

Interestingly, an initial clustering analysis indicated that cocaine exposure substantially reduced sex differences in NAc chromatin organization (**Suppl. Figure 7A**). Strikingly, when we compared the differentially accessible regions (DARs) between males and females, we found that acute cocaine exposure leads to a more than 10-fold reduction in the number of sex-specific open chromatin regions in both male-diestrus and male-proestrus comparisons (**Suppl. Figure 7B**). We then focused on the changes in chromatin occurring between the control and cocaine conditions separately in each group – diestrus, proestrus, and males (**Fig. 4B-D; Suppl. Table 4**). Initially, we found minimal overlaps in the specific cocaine-induced DARs between groups (**Figure 4B**). However, once DARs were annotated to the nearest gene, we identified a similar number of genes with cocaine-induced chromatin changes in males, diestrus females, and proestrus females, with a much higher overlap between-groups (**Figure 4C**). Interestingly, the highest number of genes with affected chromatin found in males (7563) was in line with the more consistent CPP effect found in control males at this dose, compared to control females (**Figure 1C**). Within females, proestrus mice had a lower number of genes with cocaine-induced chromatin changes (4499) compared to diestrus mice (5747), consistent with the finding that females in a high-estrogenic phase during their first cocaine exposure exhibit a lower conditioning response to cocaine (**Figure 1D**). Examining the overlapping genes between groups, we found the most substantial overlap between diestrus and males (3530), the two cocaine CPP-susceptible groups, compared to diestrus and proestrus (2260) or proestrus and males (2483, **Figure 4C**). When we performed GO and KEGG analysis on the genes with cocaine-induced DARs, we found that, overall, similar terms and pathways are affected by cocaine in each group, including synapse organization, dendritic spine organization, dopaminergic synapse, and multiple addiction-related pathways (**Figure 4D**).

In summary, neuronal chromatin organization is altered by acute cocaine exposure in males, diestrus females, and proestrus females, and is enriched near genes related to neuronal function and addiction. Similar to the effects of early-life stress on NAc gene expression, acute cocaine exposure also reduces sex differences in NAc chromatin organization.

### Acute cocaine exposure increases the accessibility of regions harboring AP1 motifs

To identify the putative upstream regulators of cocaine’s effects on neuronal chromatin, we performed motif analysis on DARs that are more accessible after acute cocaine exposure separately in each group. In all three groups, the top motif was Fos::JunB (**Figure 5A**), corresponding to the binding site for the AP1 transcription factor formed by dimers of Fos-and Jun-family proteins (46). Genes with Fos::JunB motif-containing DARs that are shared by all three groups are enriched for the Morphine addiction KEGG pathway (**Figure 5A**), consistent with the well-established role of Fos proteins in mediating addiction-related molecular phenotypes in the NAc (47). However, when we analyzed promoter DARs harboring Fos::JunB motifs, we found that diestrus females had the largest number of such DARs (689), followed by proestrus females (74), and males (37, **Suppl. Table 5**). The vast majority of Fos::JunB motif-containing DARs were sex-specific, including a male-specific DAR at the promoter of *Drd1* (**Figure 5B**), encoding the Dopamine receptor D1, a gene extensively involved in mediating cocaine’s effects (48), and a female-specific DAR at the promoter of *Gabrg1* (**Figure 5B**), encoding GABA receptor G1, a gene whose genetic variants in the human paralog have been associated with addiction risk (49). Diestrus-specific promoter DARs with AP1 sites include *Syn1* (**Figure 5B**), an X-linked gene whose expression we found to be altered following early-life stress in females (**Suppl. Figure 6B**).

**Figure 5.**
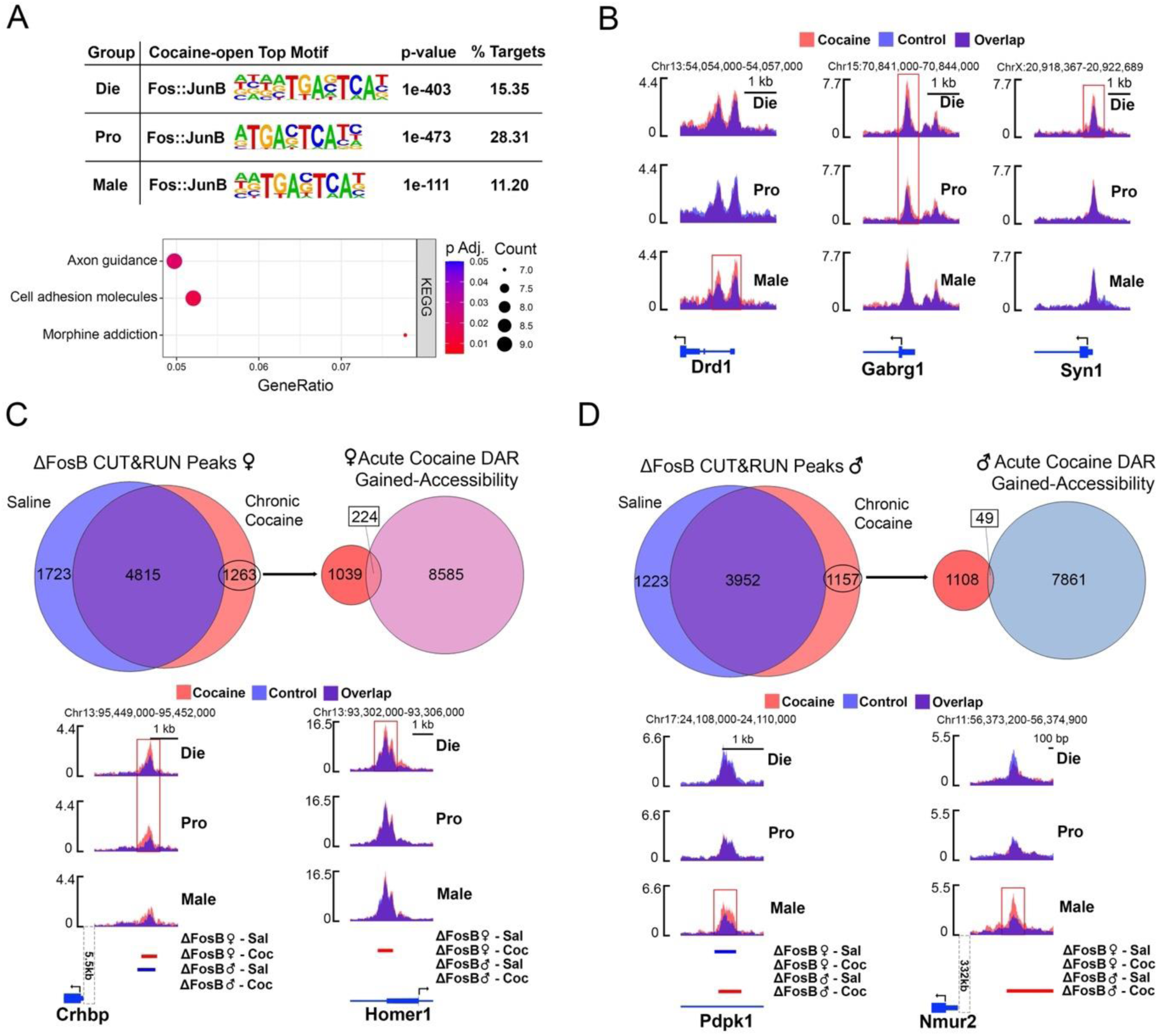
Acute cocaine exposure increases the accessibility of AP-1 motifs and ΔFosB binding sites. **(A)** Regions that gain accessibility after acute cocaine are enriched for the Fos::JunB (AP1) motif as the top motif in each group (top), and a dotplot shows the top KEGG pathways enriched in genes with cocaine-induced, AP1 motif-containing open regions in all three groups (bottom). Colors indicate adjusted *p-*values and dot size corresponds to gene count. **(B)** SparK plots of group-average normalized ATAC-seq reads (n = 2-3 replicates or 6-9 animals/group) are shown for genes with gained accessibility of Fos::JunB motifs after acute cocaine, including regions near the transcription start sites (TSSs) of *Drd1* (left), which gains accessibility only in males, *Gabrg1* (middle), which gains accessibility only in females, and *Syn1* (right), which gains accessibility only in diestrus females. Red boxes depict the differentially accessible regions (DARs). **(C)** Using CUT&RUN data (Yeh et al., 2023) of ΔFosB binding in female NAc Drd1-expressing medium spiny neurons (D1-MSNs), we focused on regions specifically bound by ΔFosB in the chronic cocaine condition (left Venn diagram) and overlapped them with regions that gain accessibility in females after acute cocaine in our ATAC-seq data (right Venn diagram). Shown below are SparK plots of group-average normalized ATAC-seq reads (n = 2-3 replicates or 6-9 animals/group) for overlapping genes including a region upstream of the TSS of *Crhbp*, which exhibits female-specific gain of accessibility and ΔFosB binding after acute and chronic cocaine, respectively (left), and a region near the TSS of *Homer1* which exhibits diestrus-specific accessibility gain after acute cocaine and female-specific ΔFosB binding after chronic cocaine (right). Red boxes depict the differentially accessible regions (DARs), bed tracks below depict ΔFosB binding in the chronic cocaine and chronic saline conditions for males and females. **(D)** Using CUT&RUN data (Yeh et al., 2023) for ΔFosB in male D1-MSNs, and as in **(C)**, we focused on regions bound by ΔFosB specifically in the chronic cocaine condition (left Venn diagram) and overlapped them with regions that gain accessibility in males after acute cocaine in our ATAC-seq data (right Venn diagram). Shown below are SparK plots of group-average normalized ATAC-seq reads (n = 2-3 replicates or 6-9 animals/group) for overlapping genes including an intronic region of *Pdpk1*, which exhibits male-specific gain of accessibility and ΔFosB binding after acute and chronic cocaine, respectively (left), and a region upstream of *Nmur2*, which exhibits male-specific gain of accessibility and ΔFosB binding after acute and chronic cocaine, respectively (right). Red boxes depict the differentially accessible regions (DARs), bed tracks below depict ΔFosB binding in the chronic cocaine and chronic saline conditions for males and females. Die, diestrus; Pro, proestrus; Male, males.

Sites harboring the Fos::JunB motif could be bound by AP1 proteins such as cFos, which is implicated in mediating the *acute* effects of cocaine (50), as well as by ΔFosB which has been extensively studied for its role in the transcriptional effects of *chronic* cocaine exposure (50–55). We, therefore, tested the overlap of our regions that gain chromatin accessibility after acute cocaine exposure with the regions bound by cFos (56) or ΔFosB (57) derived from publicly available ChIP-seq and CUT&RUN datasets, respectively. This analysis revealed greater overlap between cocaine-induced DARs and ΔFosB-bound regions compared to cFos-bound regions in all samples, and most profoundly in the diestrus group (**Suppl. Figure 8**), indicating that cocaine acutely alters chromatin accessibility in targets of ΔFosB, priming them for regulation following chronic cocaine exposure. To further evaluate this hypothesis, we performed the overlap of regions that gain accessibility after acute cocaine in our data with regions bound by ΔFosB in Drd1-expressing medium spiny neurons (D1 MSNs) specifically after chronic cocaine treatment in the CUT&RUN dataset by Yeh et al. (57). We found that 17.7% (or 224 out of 1263) of these regions, bound by ΔFosB in the chronic cocaine, but not saline condition, gain accessibility after acute cocaine exposure in females (**Figure 5C**). Of these regions, 49.6% are unique to diestrus, 25.4% are unique to proestrus, and 25% are shared between the two female groups (**Suppl. Table 5**). Examples include a region upstream of *Crhbp*, encoding Corticotropin releasing hormone-binding protein, shown to be involved in both addiction (58, 59) and the sex differences in the stress response (60). This *Crhbp* region gains accessibility after acute cocaine and gains ΔFosB binding after chronic cocaine specifically in females (**Figure 5C**). In fact, this same region loses ΔFosB binding after chronic cocaine treatment in males (**Figure 5C**). The promoter region of *Homer1*, encoding a transcription factor that plays a role in cocaine-induced synaptic plasticity (61), similarly gains ΔFosB binding after chronic cocaine only in females, but only gains chromatin accessibility after acute cocaine in diestrus females (**Figure 5C**). Notably, in males, only 4.2% (or 49 out of 1157) of regions that gain ΔFosB binding after chronic cocaine exposure overlap with a region that gains accessibility after acute cocaine exposure (**Figure 5D**). These regions include an intronic region of *Pdpk1*, encoding 3-phosphoinositide-dependent protein kinase-1, and a region upstream of *Nmur2* (**Figure 5D**), encoding a neuropeptide receptor whose activity is linked to addiction-related phenotypes (62) and whose expression in the NAc is dysregulated following repeated cocaine administration (63). Together, these data show that acute cocaine exposure sex-specifically increases chromatin accessibility in AP1 motif-containing regions that exhibit sex-specific ΔFosB binding after chronic cocaine. Notably, diestrus females had the greatest number of overlaps, and gained accessibility of ΔFosB binding sites therefore represents a putative epigenomic mechanism through which sex hormone status in females during an acute exposure to cocaine can affect susceptibility to future exposures.

### Cocaine’s effects on chromatin accessibility are sex-and estrous cycle stage-dependent

In order to explore the relationship between changes in cocaine-induced chromatin accessibility and behavioral phenotypes, we focused first on the overlapping genes with cocaine-induced DARs in diestrus and males, since these groups showed stronger cocaine-induced conditioning responses in the CPP test (**Figure 1C, D**). Enrichment analysis revealed enrichment for genes related to GABA-ergic signaling, learning, and dendritic spines (**Figure 6A**). As examples, we observed changes in the promoter regions of three genes of interest:*Sp9*, encoding a transcription factor with a role in MSN development (64), *Fgfr1*, encoding fibroblast growth factor receptor 1, whose activity is linked to addiction related phenotypes (65, 66) and *Gad2*, encoding glutamate decarboxylase 2, an enzyme which facilitates the conversion of glutamate to GABA ((67), **Figure 6A, Suppl. Table 4**).

**Figure 6.**
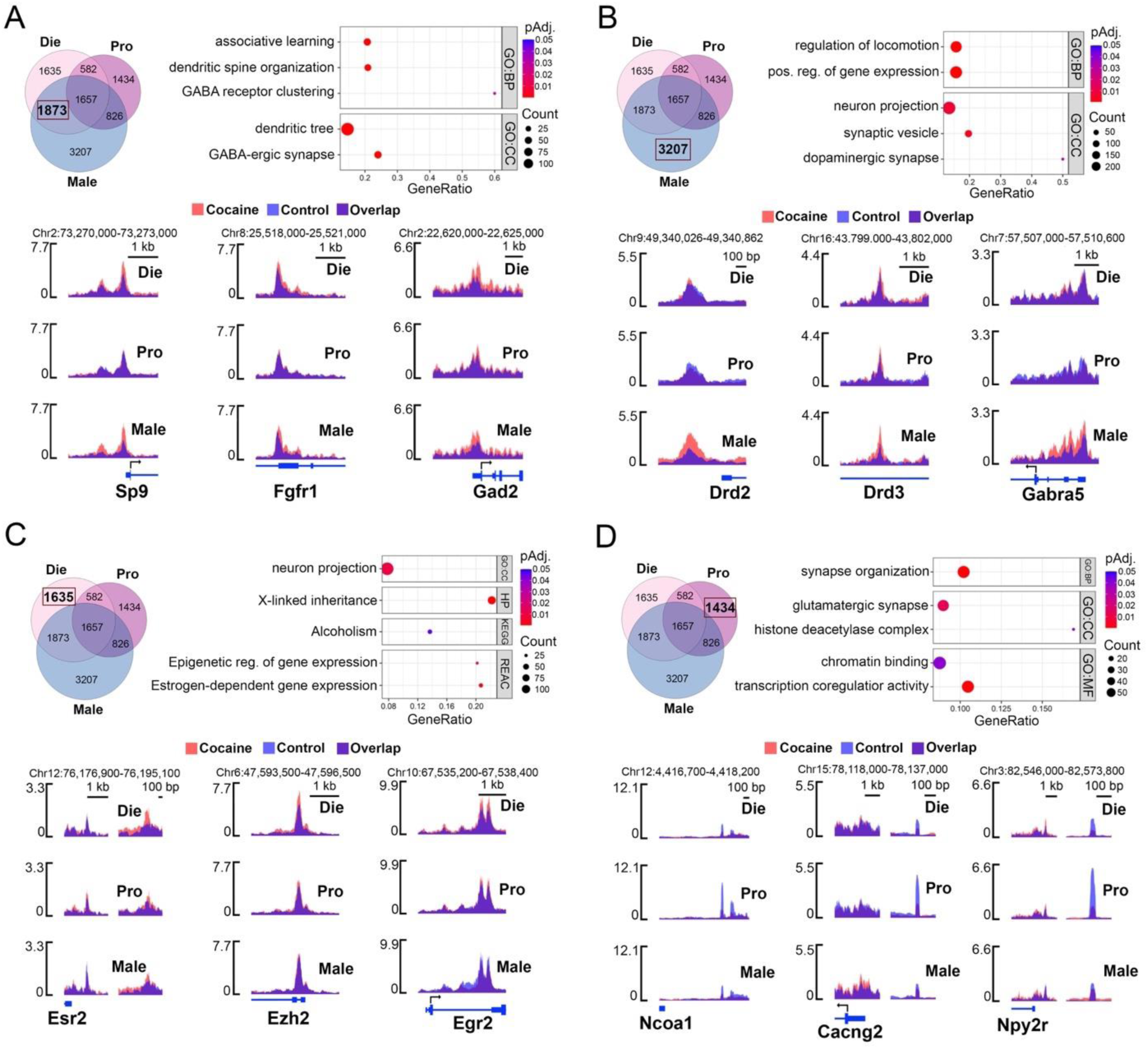
Acute cocaine alters chromatin accessibility in a sex-and estrous cycle stage-specific manner. **(A)** A dotplot (right) shows select gene ontology (GO) terms for biological process (BP) and cellular component (CC), enriched in genes annotated to cocaine-induced differentially accessible regions (DARs) shared by males and diestrus females, but not proestrus females (shown in the Venn diagram, left). SparK plots of group-average normalized ATAC-seq reads (n = 2-3 replicates or 6-9 animals/group) are shown below for male and diestrus overlapping genes including regions that gain accessibility near the transcription start sites (TSSs) of *Sp9* (left), *Fgfr1* (middle), and *Gad2* (right). **(B)** A dotplot (right) shows select BP and CC gene ontology terms enriched in male-specific genes annotated to cocaine-induced DARs (shown in the Venn diagram, left). SparK plots of group-average normalized ATAC-seq reads (n = 2-3 replicates or 6-9 animals/group) are shown below for male-specific genes including regions that gain accessibility near the TSS of *Drd2* (left), within an intron of *Drd3* (middle), and near the TSS of *Gabra5* (right). **(C)** A dotplot (right) shows select CC gene ontology terms, as well as pathways from the human protein atlas (HP), KEGG, and Reactome (REAC) databases enriched in diestrus-specific genes annotated to cocaine-induced DARs (shown in the Venn diagram, left). SparK plots of group-average normalized ATAC-seq reads (n = 2-3 replicates or 6-9 animals/group) are shown below for diestrus-specific genes including regions that gain accessibility upstream of *Esr2* (left), near the TSS of *Ezh2* (middle), and near the TSS of *Egr2* (right). **(D)** A dotplot (right) shows select BP, CC, and molecular function (MF) gene ontology terms enriched in proestrus-specific genes annotated to cocaine-induced DARs (shown in the Venn diagram, left). SparK plots of group-average normalized ATAC-seq reads (n = 2-3 replicates or 6-9 animals/group) are shown below for proestrus-specific genes including regions that lose accessibility upstream of *Ncoa1* (left), *Cacng2* (middle), and *Npy2r* (right). For all dotplots, colors indicate adjusted *p-*values and dot size corresponds to gene count. Die (light pink), diestrus; Pro (purple), proestrus; Male (blue), males.

While there was a substantial overlap in genes with cocaine DARs between diestrus females and males, it is also plausible that the two groups reach a “susceptible state” through independent mechanisms. We therefore separately focused on genes with cocaine-induced DARs specific to each group. Male specific genes were enriched for terms including positive regulation of gene expression, synaptic vesicle, and, interestingly, dopaminergic synapse (**Figure 6B**). While the Dopaminergic synapse KEGG pathway was commonly enriched in all three groups (**Figure 4D**), a subset of genes involved in the dopaminergic synapse undergo chromatin accessibility changes in males only. Examples include genes encoding the dopamine receptors *Drd1* (**Figure 5B**), *Drd2*, and *Drd3* (**Figure 6B, Suppl. Table 4**). We also identified genes involved in GABA-ergic signaling with male specific cocaine DARs, including *Gabra5*, encoding a GABA receptor whose activity in the human male NAc is associated with addiction ((68), **Figure 6B, Suppl. Table 4**).

Enrichment analysis of diestrus-specific genes revealed pathways including X-linked inheritance and Estrogen-dependent gene expression (**Figure 6C**), indicating that both sex hormone signaling and the X chromosome are important features of the diestrus female response to acute cocaine exposure. For example, cocaine exposure increases chromatin accessibility in a region upstream of the *Esr2* locus, encoding ERβ, specifically in diestrus females (**Figure 6C, Suppl. Table 4**). Notably, gene expression of the ERβ signaling pathway was altered in females by early-life stress (**Figure 3A-C, Suppl. Table 3**), and therefore represents a molecular feature shared by two cocaine susceptible groups:diestrus females and females exposed to early-life stress. We also observed diestrus-specific DARs at the promoters of *Ezh2*, encoding a histone methyltransferase, and *Egr2*, encoding early growth response 2, an immediate early gene whose induction in the NAc by cocaine is required for establishing cocaine CPP ((69), **Figure 6C, Suppl. Table 4**).

In summary, males and diestrus females exhibit similar changes in cocaine accessibility in genes related to GABA-ergic function, but also have divergent responses, with male-specific DARs observed at dopamine receptor genes and diestrus-specific DARs observed at genes related to estrogen signaling. To better understand why high-estrogenic females are less susceptible to the conditioning effects of cocaine (**Figure 1D**), we also performed enrichment analysis on genes with proestrus-specific DARs (**Figure 6D**). We identified terms related to gene and chromatin regulation, including histone deacetylase complex, chromatin binding, and transcription coregulator activity (**Figure 6D**). One example is *Ncoa1*, a transcriptional coregulator involved in mediating the transcriptional effects of steroid hormones (70) in which a region of chromatin upstream of the TSS becomes less accessible in proestrus females after cocaine exposure (**Figure 6D, Suppl. Table 4**). Notably, this region of chromatin is more accessible in proestrus controls compared to diestrus and male controls, indicating chromatin would typically gain accessibility in this region as an element of a proestrus-specific chromatin state. Then, after cocaine exposure, this region becomes more similar to diestrus and males in terms of chromatin accessibility. We observed this same pattern at genes including *Cacng2*, a calcium channel involved in regulating AMPA receptor density following cocaine-sensitization (71) and *Npy2r*, a neuropeptide receptors whose polymorphisms in humans are associated with cocaine addiction ((72), **Figure 6D, Suppl. Table 4**). Interestingly, when we overlapped proestrus cocaine-induced DARs that are less accessible after cocaine with DARs that are more accessible in proestrus vs. diestrus controls, we found enrichment for genes related to dendritic spines, synapse organization, and behavior (**Suppl. Figure 9**). This interruption of proestrus DARs by cocaine exposure indicates a competition between cocaine-and proestrus state-induced chromatin regulation, which may be protective for proestrus females by limiting the effects of cocaine on neuronal chromatin organization.

### Cocaine disproportionately leads to chromatin opening on the X chromosome in diestrus females

Corroborating the notion that the proestrus phase is protected from the effects of cocaine on chromatin, we found that, in addition to having less cocaine-induced DARs than diestrus, proestrus DARs are mostly regions that are less accessible after cocaine (55.7%). By contrast, in diestrus, the majority of DARs are regions more accessible after cocaine (66.8%, **Figure 7A**). This pattern was most striking in gene promoters; in these regions, 83% of proestrus-specific DARs lose accessibility and 88% of diestrus-specific DARs gain accessibility after acute cocaine exposure (**Suppl. Figure 10A**). Moreover, gene promoters that become less accessible in proestrus after cocaine are enriched for terms and pathways related to neuronal function, addiction, and, importantly, estrogen signaling (**Suppl. Figure 10B**). To explore the preponderance of gained-accessible regions in diestrus, and given the enrichment for the X-linked inheritance term in diestrus specific genes (**Figure 6C**), we assessed whether this ratio of open:closed DARs varied by chromosome. Indeed, we found that this ratio was particularly skewed on the X-chromosome in diestrus females where practically all of the chromatin changes were characterized by more accessible chromatin, which was not the case in proestrus females (**Figure 7B**). Surprisingly, X-linked genes whose chromatin was impacted by cocaine in diestrus females included the same X-linked lncRNAs whose expression was disrupted by early-life stress in females (**Figure 3E, Suppl. Table 2**), including Xist, Jpx, and Firre (**Figure 7C, Suppl. Table 4**). In each case, chromatin was more accessible near the TSS after cocaine in diestrus females only. In fact, we observed the same pattern in autosomal genes involved in X-inactivation whose expression is altered by early-life stress in females (**Suppl. Figure 6A**, **Suppl. Table 2**), specifically *Ythdc1*, *Lrif1,* and *Smchd1* all exhibit increased chromatin accessibility near the TSS after cocaine in diestrus females (**Suppl. Figure 11, Suppl. Table 4**). In addition, we found that X-linked genes with cocaine-induced DARs in diestrus were enriched for synapse related terms and pathways (**Figure 7D**). Enrichment analysis of these same genes specifically using the Reactome pathway set revealed enrichment of three neuronal pathways:Neuronal system, Glutamatergic synapse, and Protein interactions at synapse, including genes such as *Syn1* and *Gria3* known to be critical for synaptic plasticity (**Figure 7E**). In summary, cocaine exposure affects chromatin regulation of the X-chromosome preferentially in diestrus females, affecting genes involved in dosage compensation as well as synaptic function which can, in part, explain the susceptibility of diestrus females to the conditioning effects of cocaine.

**Figure 7.**
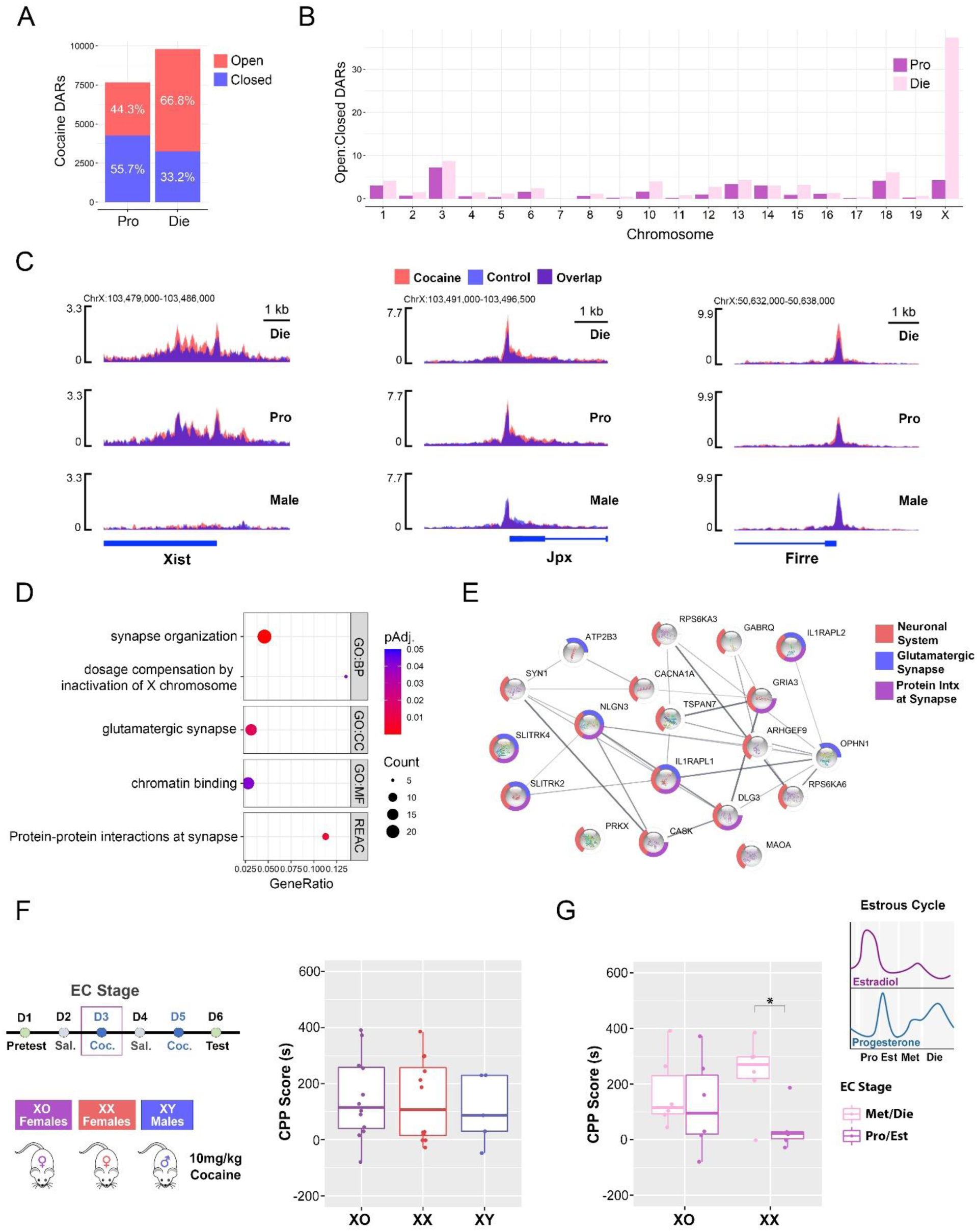
The role of the X chromosome in the cocaine response is estrous cycle-dependent. **(A)** Among cocaine-induced differentially accessible reagions (DARs), more chromatin regions become open (red) than closed (blue) in diestrus, while it is opposite in proestrus. **(B)** Across chromosomes, the ratio of Open:Closed cocaine-induced DARs is particularly skewed on the X chromosome for diestrus, but not proestrus. **(C)** SparK plots of group-average normalized ATAC-seq reads (n = 2-3 replicates or 6-9 animals/group) are shown for the transcription start sites of the X-linked long non-coding RNAs *Xist* (left), *Jpx* (middle), and *Firre* (right), which all gain accessibility after cocaine specifically in diestrus females. **(D)** A dotplot shows select gene ontology (GO) terms for biological process (BP), cellular component (CC), and molecular function (MF), as well as Reactome (REAC) pathways enriched in X-linked genes with cocaine-induced DARs in diestrus. Colors indicate adjusted *p-*values and dot size corresponds to gene count. **(E)** Reactome pathways enriched in X-linked genes with cocaine-induced DARs in diestrus were visualized using String, with nodes corresponding to genes color-coded by pathway and edges corresponding to protein-protein interactions between gene products. **(F)** The CPP test was performed, following the same timeline as previous experiments (top-left), with 39,XO females, XX females, and XY males of the same genetic background using a 10 mg/kg cocaine dose for conditioning (bottom-left). CPP scores were analyzed in all 3 groups using one-way ANOVA (n = 12 females/group; n = 5 males). **(G)** The CPP score analysis was repeated in female mice considering their estrous cycle stage on the first day of cocaine exposure. This analysis was done with group and estrogen status as factors (Two-way ANOVA, n = 6/stage/group). *p < 0.05; post-hoc Welch two-sample T-test. Box plots (box, 1st–3rd quartile; horizontal line, median; whiskers, 1.5× IQR). Pro, proestrus; Est, estrus; Met, metestrus; Die, diestrus; Male, males.

### The inactive X chromosome is required for the estrous cycle’s effect on cocaine-induced CPP

To further address the importance of the X chromosome on cocaine’s effect in female mice, we performed another CPP experiment comparing female mice with only one X chromosome (39, X0), wildtype female mice (40, XX) and male mice (40, XY) of the same genetic background (**Figure 7F**). While animals in each group, on average, developed a preference for the cocaine-paired compartment, we did not detect a difference in CPP scores between the three groups (F(_2, 26_)= 0.155, p=0.86; one-way ANOVA; **Figure 7F**). Next, we tested the effect of ovarian hormone-status on the day of first cocaine exposure on CPP scores and found a significant effect of the estrous cycle (F_(1, 20)_= 5.235, p=0.03; two-way ANOVA; **Figure 7G**), with females in a low-estrogenic phase (Met/Die) on the first day of cocaine exposure exhibiting higher CPP scores than females in the high-estrogenic phase (Pro/Est), as we showed previously (**Figure 1D**). However, when we followed up on this result with a post-hoc test, we found that this effect could only be detected in the XX group (t(_5_)=3.23, p=0.01; Welch Two Sample t-test), while it was absent in the 39,XO group (t(_5_)=0.51, p=0.62; Welch Two Sample t-test), indicating the estrous cycle’s effect on cocaine CPP requires the presence of two X chromosomes in females.

## Discussion

In this study, we show that two sex-specific risk factors for CUD, early-life stress and ovarian hormone status, induce sex-specific responses to cocaine in mice, at both the behavioral and molecular level. Although cocaine can induce CPP in both males and females, the effect is dose-dependent and more consistent in males, while it varies with the estrous cycle stage and is more strongly affected by early-life stress in females. Interestingly, we observe both shared and distinct effects of early-life stress and ovarian hormone status on transcriptional regulation in the key reward area NAc and we link them to sex-specific responses to cocaine.

In control animals, we show a more consistent cocaine-induced conditioning effect with 10 mg/kg dose in males than in females. The result in males was expected since the dose was selected based on the previous research performed predominantly in males (19). The result in females, on the other hand, requires a more careful consideration. The previous study by Satta et al. also reported less consistent cocaine CPP effect in females (4) but was performed with a lower (5 mg/kg) dose of cocaine. Varied response to cocaine in females is consistent with the idea that the cocaine effect may vary with the estrous cycle stage (2, 5). However, the well-established finding is that estrogen facilitates cocaine-induced dopamine release and that high-estrogenic phases (typically proestrus or estrus) are associated with an increase in cocaine self-administration (5, 73).

Here, however, we report that the high-estrogenic phase of the cycle associated with the first exposure to cocaine results in a weaker conditioning response to cocaine. This response may reflect the supposedly protective effect of estrogen against cocaine-induced synaptic plasticity in the NAc, independent of the acute, dopamine-related reinforcing effect of estrogen. On the other hand, the low-estrogenic phase in females seems to be the most susceptible state to facilitate cocaine-induced CPP, as observed at both 10 mg/kg as well as at 2.5 mg/kg cocaine that is insufficient to induce CPP in males. One plausible reason for this can be the negative affective state in mice during diestrus in which we and others showed consistently higher anxiety indices compared to the high-estrogenic, proestrus phase (10, 15, 74). Consistent with this, earlier studies showed that the first exposure to cocaine has acute anxiolytic effects (75) which may facilitate the association of cocaine exposure with improved affective state in mice. This is further consistent with the evidence in humans that, unlike men who seek drugs to engage in risk-taking behaviors, women initiate drug use as a coping strategy to deal with anxiety, depression, and feelings of isolation (73).

Thus, the overall effect of cocaine on CPP is more complex in females than in males and we observe both protective and predisposing effects of ovarian hormones, depending on the estrous cycle phase. Interestingly, early-life stress erases the effect of the cycle and make all females susceptible to cocaine CPP, along with males at their otherwise subthreshold dose. This effect of early-life stress, again, partially mimics the effect of early stress seen in humans, where women are more strongly affected than men (73). However, both sexes show susceptibility to early-life stress enhancing addiction risk later in life (76).

We performed a gene expression study to address the underlying molecular mechanism for the sex-specific effect of early-life stress on cocaine-induced behavioral phenotypes. Two earlier studies characterized gene expression changes in the NAc, the key brain reward area, after early-life stress (11, 12). These researchers used different paradigms from ours, including both different stressors (limited bedding with or without MS) as well as different developmental time points (P10-17 or P2-10), so it is unsurprising that our gene expression data differ as well. However, we do observe shared genes relevant to neurodevelopment and, importantly, all paradigms show a stronger effect of early-life stress on NAc gene expression in females than in males (11, 12), which is most evident in our study. In addition to the female-biased response, we also observe an overall decreased sex difference in the NAc gene expression following early-life stress, similar to the previously revealed effect of adolescent social stress on the NAc transcriptome (77), and mimicking the pattern found in our behavioral CPP data.

There are two important findings that can help explain the female-biased effect of early-life stress on NAc gene expression and cocaine-induced CPP. First, we see the enrichment of genes involved in estrogen signaling in response to early-life stress in females (but not in males). Specifically, ERβ in the NAc was previously shown to regulate the development of cocaine CPP in female mice (4). Second, and even more intriguingly, we find an overwhelming response of X chromosome-linked genes to early-life stress in females, with a particularly intriguing increase in the expression of genes involved in the female-specific phenomenon of X chromosome inactivation. Both of these mechanisms, acting separately or in concert, can induce a female-specific response to early-life stress of relevance to addiction. Specifically, the majority of down-regulated genes in females are involved in synaptic function pointing toward altered regulation of synaptic plasticity after early-life stress. On the contrary, upregulated genes show enrichment of genes involved in transcriptional and chromatin regulation, known to be important in the acute and chronic effects of cocaine.

Early-life stress effectively nullifies the effect of the ovarian hormone status on cocaine-induced CPP in females, however, as mentioned above, these hormones have a critical effect on cocaine CPP in females under basal conditions. Thus, we further addressed the molecular mechanisms through which ovarian hormones contribute to a female-specific response to cocaine. Among transcriptional mechanisms, chromatin regulation in the NAc was the major candidate mechanism. As previously mentioned, chromatin regulation in the NAc is strongly implicated in cocaine-induced adaptations underlying addiction-related behaviors, although the evidence comes from studies almost exclusively performed in males (9). However, we previously reported sex-and estrous cycle-dependent chromatin regulation in the related brain area, the ventral hippocampus (10). Here, we importantly show that acute cocaine treatment induces sex-and estrous cycle-dependent changes in neuronal chromatin within the NAc, providing a plausible molecular substrate to explain sex-specific responses to cocaine. Interestingly, we again observe that cocaine, similar to early-life stress, reduces sex differences in NAc neuronal chromatin organization that exist under basal conditions.

While a lot of genes and pathways that are affected by cocaine are shared between males and females across the cycle, there are several sex-and estrous cycle-specific molecular features in our ATAC-seq data that reveal candidate sex-specific transcriptional mechanisms underlying cocaine-induced responses. First, we show the enrichment of AP1 binding motifs within chromatin regions with increased accessibility in response to cocaine. While cFos and other Fos proteins have been strongly implicated in the acute response to cocaine (50), to our knowledge, this is the first report that shows the genome-wide enrichment of AP1 sites within the cocaine-induced changes in NAc chromatin organization. Interestingly, AP1 sites were overwhelmingly enriched within the promoter regions during diestrus, the low-estrogenic state that represents the most susceptible state for the development of cocaine preference. While the chromatin changes that we observed are induced by acute cocaine treatment, we have an indication that they are relevant to chronic cocaine exposure as well. We show that AP1 binding sites that gained chromatin accessibility in response to acute cocaine strongly overlapped with ΔFosB binding, the Fos protein involved in the transcriptional effects of chronic cocaine exposure (50–55). And, considering that this overlap was most profound in the diestrus group, we propose that the low-estrogenic state in females, in response to acute cocaine exposure, primes chromatin for later binding of ΔFosB and other molecular adaptations following chronic cocaine exposure. Interestingly, a recent study showed that estrogen withdrawal increases ΔFosB expression in the NAc (78), consistent with our hypothesis that a physiological drop in estrogen (in diestrus) can “prime” chromatin for cocaine-induced long-term plasticity in the NAc.

Interestingly, we also observed more overlaps in chromatin changes between diestrus and males, the two groups “more susceptible” to cocaine CPP, compared to proestrus. These overlaps are certainly interesting as they include addiction-related biological pathways such as dendritic spine density, learning, and GABAergic synapse. However, equally interesting is the possibility that these two groups may achieve a similar behavioral phenotype through different mechanisms. For instance, the enrichment of dopaminergic synapse-related genes has been the major finding reported by previous studies, again largely performed in males (79, 80). While our data confirm that dopaminergic pathway is shared across sex we still see important male-specific chromatin changes in genes encoding dopamine receptors, which are likely important for sex-specific cocaine-induced responses and could have not been anticipated by earlier, male-focused studies. Interestingly, diestrus-specific cocaine-induced DARs are enriched in X-linked and estrogen signaling-related genes, which includes diestrus-specific chromatin opening within the *Esr2* gene encoding ERβ. These two mechanisms, related to ERβ signaling and X chromosome-linked gene expression, are shared between the two susceptible female groups, control diestrus and early-life stress-exposed females, revealing female-specific transcriptional mechanisms underlying cocaine induced preference.

We would specifically like to emphasize a coincidence of the increased expression and increased chromatin accessibility of the X-linked lncRNAs (Xist, Jpx, Firre) in females after early-life stress and after cocaine exposure in diestrus, respectively. Considering the important roles that these lncRNAs play in maintaining dosage compensation (29–31, 33), our findings indicate a global disruption in the regulation of the inactive X chromosome in the two cocaine-susceptible female groups. In general, dysregulation of the X chromosome can contribute to the altered neuronal function underlying the early-life stress-induced and diestrus-driven susceptibility to cocaine, given the overrepresentation of genes related to brain development and function on the X chromosome (81). Accordingly, the role of X chromosome in cocaine response was previously addressed by the four core genotype model and it was reported that the XX genotype, regardless of sex hormones, affects cocaine vulnerability (82). Here, though, we used 39,XO mice as a model to reveal the functional role of the inactive X chromosome in response to cocaine within the naturally-cycling condition in female mice. Unlike humans with Turner syndrome (45, X0) whose ovaries are either missing or do not function properly, 39,XO mice (39,X0) have functional ovaries and undergo the estrous cycle (83). Using the 39,XO model, thus, we show that female mice that have only one X chromosome are not able to respond to the protective effect of physiologically high estrogen levels. And, similar to the effect of early-life stress or physiological estrogen withdrawal, lack of the inactive X chromosome leads to enhanced susceptibility to cocaine-induced conditioning response in females.

In addition to female-specific susceptibility, we would also like to address the molecular mechanisms that may underlie the “protective effect” of estrogen against cocaine-induced synaptic plasticity in the NAc. Interestingly, we see a preferential closing of chromatin in proestrus in response to acute cocaine exposure, unlike the largely increased chromatin accessibility we observed in the susceptible, diestrus group. This interruption of proestrus-specific open chromatin regions by cocaine exposure indicates a competition between cocaine-and estrogen-induced chromatin regulation, which may be protective for proestrus females by limiting the effects of cocaine on neuronal chromatin organization. However, we would like to reiterate that previous studies have shown estrogen-induced enhancement of cocaine’s reinforcing effects (73). Considering that this effect may be mediated via fast estrogen signaling (2), we do not believe that it is necessarily inconsistent with estrogen’s protective effect that we observed on cocaine-induced chromatin plasticity, which is slower and has more long-term consequences. These data are also consistent with our finding that the low (but not high) estrogenic state in females interacts with acute cocaine exposure and primes chromatin for later binding of ΔFosB and other molecular adaptations associated with chronic cocaine exposure.

Substance use disorders involve multiple components from motivational aspects and initiation of drug use to adaptive changes in the brain that underlie chronic drug use and addiction. Our study highlights that cocaine use in women and other menstruating individuals may involve complex effects of ovarian hormones that interact with internal factors (such as negative affective state) and external risk factors such as stress and, for the first time, we reveal the molecular substrates through which female-specific factors can induce stronger and longer-lasting, sex-specific responses to cocaine. Our findings offer a molecular framework to understand sex-specific mechanisms underlying cocaine use disorder, opening new avenues for future sex-and gender-informed treatments for CUS.

## Methods

### Animals

For early-life stress experiments, five-week-old male (n=18) and four-week-old female (n=36) C57BL/6J mice were obtained from Jackson Laboratories. After two weeks of habituation period, mating pairs (n=18) were formed consisting of one male and two female mice and were separately housed. After mating, n=34 of the subsequent litters were used in this study. Litters were counted at P0, and litters were randomly assigned to either Con or MS groups (n=17 litters/group). Litters were weaned at P28 and housed in cages with mice of the same sex and group (n=5 per cage). For acute cocaine treatment (ATAC-seq) experiments, four-week-old male (n=18) and female (n=48) C57BL/6J mice were obtained from Jackson Laboratories. Animals underwent estrous cycle monitoring for approximately two weeks (see *Estrous Cycle*), prior to acute cocaine treatment at P52-54. For the X-chromosome experiment, 39,XO females (n=12), XX females (n=12), and XY males (n=5) of the same genetic background (C57Bl/6J x CBA/CaGnLeJ, Strain #036414) were obtained from Jackson Laboratories. The genotype of animals was confirmed in-house using the established qPCR assay (Jackson Laboratories Protocol #35972) performed on DNA isolated from tail clippings. All animals used in the study were housed in same-sex cages (n=3-5/cage), were kept on a 12h:12h light:dark cycle, were given *ad libitum* access to food and water, and were allowed to habituate to the facility for at least two weeks prior to experiments. All animal procedures were approved by the Institutional Animal Care and Use Committee at Fordham University.

### Early-life stress

To administer early-life stress, a well-established maternal separation protocol was used (14–16). Litters chosen for early-life stress (MS group) were separated from the dams for three hours each day from P1-P14. Control litters (Con group) did not undergo any manipulations during this time. The separation window was changed each day to ensure that the stress was unpredictable. During separation, the dam was placed in a temporary cage with food and water. Dams were exposed to restraint stress (20min) or forced swim stress (2min) at a random time during the three-hour separation window in order to reduce compensatory maternal care following separation, which has been reported to occur (17). We confirmed that our unpredictable stress protocol disrupts maternal behavior for at least one hour after the mother is returned to the home cage (**Figure 1B**).

### Maternal observations

The procedure for assessing maternal behavior was performed as established previously (15, 18, 84). Maternal behavior was scored from Day 1 through Day 6 postpartum in both Con and MS dams following the separation of MS dams from their pups. Every 3 minutes during the one-hour observation period, the maternal behavior during the time of inspection was recorded for each cage. Catalogued behaviors included nursing, arched-back nursing, licking and grooming, in contact with pups (not nursing), out of nest, eating, drinking, and nest building.

### Cocaine-induced Conditioned Place Preference (CPP).

From P54-P60, Con (n=50 females, 30 males) and MS (n=30 females, 30 males) animals underwent cocaine-induced conditioned place preference (CPP) testing. At this point mice were further grouped by cage into a low dose (LD) group, receiving a subthreshold dose of 2.5 mg/kg i.p. cocaine on conditioning days, and high dose (HD) group, receiving a standard dose of 10 mg/kg i.p. cocaine. For the X chromosome experiment, females (n=12/group) and males (n=5) underwent the same CPP test with only the standard 10 mg/kg dose of cocaine. Animals in this experiment were 11-13 weeks old at the time of testing, and the smaller number of males were included as a reference group to provide a comparison of the CPP phenotype to the C57Bl/6J cohort of animals across sex. Our CPP apparatus is custom made (Maze Engineers; Skokie, IL) and consists of two conditioning chambers separated by a corridor with sliding walls. The conditioning chambers were designed to contain both visual and tactile cues, with one chamber containing striped walls and metal bars on the floor (Compartment 1), and the other containing grey walls and a plastic-holed floor (Compartment 2, **Suppl. Figure 9A**). A pilot experiment with a separate cohort of animals was used to confirm that neither conditioning chamber was generally preferable to the mice in the absence of pairing stimuli (**Suppl. Figure 9B**). The testing paradigm involved a pretest day, followed by 4 conditioning days (alternating between saline and cocaine), and finishing with a test day (**Figure 1**). During both pretest and test days, the walls were removed so that the mice could freely roam either chamber in the apparatus. During the conditioning days, mice were administered either cocaine or saline and the walls were inserted into the apparatus to confine the mice to one compartment. We used an unbiased CPP protocol in which, following injection of the mice with cocaine (2.5 or 10 mg/kg, i.p.), half were confined to Compartment 1 and the other half were confined in Compartment 2. Cocaine preference was established by calculating the CPP score (85) which was the time the animal spent in the cocaine-paired compartment on test day minus the time spent in the cocaine-paired compartment on pretest day. This score accounts for any baseline compartment preference or aversion in individual animals.

### Estrous cycle staging

We assessed the estrous cycle stage via vaginal smear cytology following a previously established protocol (10). Briefly, female mice were assigned to four possible stages of the cycle (proestrus, estrus, metestrus, and diestrus) based on cellular composition in vaginal smears. Vaginal lavage smears were taken by gently aspirating distilled water into the vaginal opening with a sterile pipette and applying the smear onto a microscope slide. After drying, slides were stained with 0.1% crystal violet solution and visualized under a light microscope. The proestrus phase is marked by predominantly nucleated epithelial cells, while estrus is characterized by mostly cornified epithelial cells. Metestrus and diestrus phases are characterized by cornified and nucleated epithelial cells, as well as leukocytes, with diestrus smears containing a far greater number of leukocytes than metestrus. For the early-life stress paradigm, the vaginal smears were obtained each day during the 6-day period of the CPP test, following the pretest, conditioning, or test session. For these experiments, the statistical analysis of behavioral data was based on the classification of females into two groups:Pro/Est consisted of animals in the proestrus or estrus phases, and Met/Die consisted of animals in the metestrus or diestrus phases. For these classifications, females were grouped based on their estrous cycle stage on the first day of cocaine exposure due to the tendency of female rodents to experience disruption of the estrous cycle following cocaine administration (86). For the ATAC-seq experiments, the estrous cycle of female animals was assessed in a more comprehensive way which included daily vaginal smears for approximately two weeks (or three cycles) to establish a predictive cycling pattern for each animal. This allowed us to predict the estrous cycle stage of female animals and ensure an adequate number of proestrus (high estrogen, low progesterone) and diestrus (low estrogen, high progesterone) females on the day of cocaine administration. Post-mortem vaginal smears were taken from these animals to confirm their estrous cycle stage.

### Drugs

Cocaine-HCl was purchased from Fagron (800006) under a Controlled Substance Schedule II license granted by the United States Drug Enforcement Agency (RK0524238) and New York State Department of Health (04C0098).

### Acute Cocaine Administration

To determine the sex-specific and estrous cycle-dependent effects of acute cocaine exposure on neuronal chromatin accessibility, groups consisting of proestrus females, diestrus females, and males aged P52-54 (aligning with the age of CPP-tested mice) were given an acute dose of 10 mg/kg i.p. cocaine (100 µl of solution containing 2.5 mg/ml of Cocaine-HCl dissolved in ddH_2_O) and sacrificed by cervical dislocation 1 hr later, while controls received no cocaine and served as the 0 hr baseline (n=9/group/time-point). Whole brains were removed, bilateral NAc were dissected immediately and snap frozen in liquid nitrogen.

### RNA-seq

At P50, just prior to the onset of CPP behavioral testing, a subset of male and female MS and Con mice (n=6/group/sex) were sacrificed by cervical dislocation, brains were removed and snap frozen in hexane. The estrous cycle of female animals was determined via vaginal smear cytology, and an equal proportion of high (n=3/group; proestrus or proestrus-estrus transition) and low (n=3/group; estrus-metestrus transition, metestrus, or diestrus) estrogenic females were included. Bilateral NAc were dissected inside a cryostat using a brain matrix and 1.5 mm biopsy tissue punch. RNA was isolated from the NAc using the RNeasy Micro Kit (Qiagen). RNA quality was assessed using the Fragment Analyzer (Agilent) and all RNA samples had an RQN of ≥ 9.0. RNA samples were quantified using the Qubit RNA High-Sensitivity assay (Thermo Fisher Scientific). cDNA libraries were prepared from RNA samples using the RNA HyperPrep Kit with RiboErase (KAPA Biosystems). First, rRNA was depleted from the RNA samples by hybridizing complementary DNA oligonucleotides to rRNA followed by treatment with RNase H and DNase. The efficiency of rRNA depletion was confirmed using qRT-PCR targeting the 28S rRNA transcript before and after RiboErase treatment (Primer sequences:Forward – 5’-CCCATATCCGCAGCAGGTC-3’, Reverse – 5’-CCAGCCCTTAGAGCCAATCC-3’). Following rRNA depletion, RNA was fragmented at 94°C for 5 minutes in the presence of Mg^2+^ and first and second cDNA strands were synthesized, followed by A-tailing. Single-indexed adaptors (KAPA) were ligated and libraries were amplified by PCR. After bead-based purification, library quality was determined with the Fragment Analyzer (Agilent), and quantification was performed using the Qubit High-Sensitivity DNA assay (Thermo Fisher Scientific). 100 bp, paired-end sequencing was performed on the Nova-Seq 6000 instrument at the New York Genome Center.

### RNA-seq Analysis

Sequences obtained from the RNA-seq experiment were adapter trimmed and aligned using Star (87) to the mouse reference genome (mm10 including small contigs) using the GENCODE M15 gtf file. Following alignment, differential gene expression analysis was performed using DESeq2 (88), and significant DEGs were considered as P_adj_ < 0.10 using DESeq2’s default correction for multiple testing (Benjamini-Hochberg). Differential expression between MS and Con groups was determined for males and females separately. For the female analysis, estrous cycle status of females was incorporated into the model. A Group*Sex interaction analysis was also performed in DEseq2 to detect genes for which the change in expression between MS and Con groups was unequal between males and females. Gene coexpression clustering analysis across the four experimental conditions was performed using Clust (22). Motif analysis on genes identified in Clust clusters was performed using Homer (89). Gene list enrichment analyses were performed using gProfiler (90). Enrichment analysis was also performed using GSEA (27) with a ranked gene list and results were plotted using EnrichmentMap (91), AutoAnnotate, and GeneMANIA apps (92) in Cytoscape (93). Aligned RNA-seq read data was plotted using SparK (94). Read numbers and quality control metrics for RNA-seq libraries are shown in **Supplementary Table 6**.

### FANS

For the ATAC-seq experiments, the purification of neuronal nuclei from bulk frozen brain tissue was carried out using a protocol we previously established (95). Dissected bilateral NAc tissue was pooled from 3 animals of the same group (forming 3 biological replicates per group, see *Acute Cocaine Administration*) and frozen in liquid nitrogen. Tissue was later homogenized in a tissue douncer. Nuclei were extracted by ultracentrifugaton through a gradient of sucrose for 1hr at 24,400rpm in a 4°C Beckman ultracentrifuge. Following centrifugation through the sucrose gradient, the nuclei-containing pellet was resuspended in DPBS and incubated with a monoclonal mouse antibody targeting NeuN, a nuclear protein expressed in neurons, conjugated with the AlexaFluor-488 fluorophore (1:1000; Millipore, MAB 377X). Nuclei were also incubated with DAPI (1:1000; ThermoFisher Scientific, 62248) prior to sorting and then filtered through a 35-µm cell strainer.

Nuclei samples were sorted using the FACSAria instrument (BD Sciences) at the Albert Einstein College of Medicine Flow Cytometry Core Facility. In order to determine the gating strategy (**Suppl. Figure 13**), the following three controls were used:1. An isotype control incubated with IgG1-AlexaFluor-488 and DAPI, 2. A DAPI-only control, and 3. A NeuN-AlexaFluor-488-only control. Gates were adjusted based on events collected from control samples in order to select single nuclei from cellular debris and clumped nuclei, and to separate the NeuN+ (neuronal) and NeuN-(non-neuronal) population. From each sample we collected 68,000-75,000 NeuN+ nuclei in BSA-coated tubes containing 200µl DPBS.

### ATAC-seq

Following FANS, neuronal nuclei from the NAc were centrifuged at 2900 x g for 10 minutes at 4°C. Transposition of chromatin for ATAC-seq was then performed as described previously (10, 96, 97). The supernatant was removed after centrifugation and the nuclei pellet was resuspended in 50µL of a transposase reaction mix which contained 2.5µL Tn5 Transposase enzyme and 25µL of 2xTD reaction buffer (Nextera DNA Library Preparation Kit). Transposition occurred at 37°C for 30 minutes, after which the DNA was purified from the samples using the MinElute PCR Purification Kit (Qiagen). A PCR reaction with the following components was used to index and amplify the transposed DNA:10µL DNA, 5µL PCR Primer Cocktail (Illumina), 25µL NEBNext High-Fidelity 2x PCR Master Mix (New England Biolabs), and 5µL of Nextera i5 and i7 indexed amplification primers (Illumina). The cycling conditions were as follows:72°C for 5 minutes, 98°C for 30 seconds, then 5 cycles of 98°C for 10 seconds, 63°C for 30 seconds, and 72°C for 1 minute. After 5 PCR cycles, a 20-cycle qPCR reaction was performed with 5µL of PCR-amplified libraries to determine the optimal number of PCR cycles needed to achieve appropriate library concentration without losing library complexity (typically 4-5 additional cycles). Following PCR, libraries were purified with the MinElute PCR Purification kit (Qiagen). Library quality was determined with the Bioanalyzer High-Sensitivity DNA assay (Agilent), and quantification was performed with the Qubit High-Sensitivity DNA assay (Life Technologies) and by qPCR (KAPA Biosystems). 100bp, paired-end sequencing was performed on the Nova-Seq 6000 instrument at the New York Genome Center.

### ATAC-seq Analysis

Following sequencing, adapters were trimmed and sequences were aligned to the mouse reference genome (mm10 including small contigs) using BWA-MEM (98). Peak-calling was then performed with MACS2, as reported previously (97). After peak calling, peaks belonging to the Y chromosome, mitochondrial DNA, or blacklisted regions, as well as peaks not shared by at least two replicates were removed from the analysis. The featureCounts function of the Rsubreads package (99) was used to count the reads in each peak across replicates. The resulting count matrix was analyzed using DESeq2 (88) to determine differential chromatin accessibility between cocaine and control groups separately for males, diestrus females, and proestrus females. Significant peaks were annotated to the nearest gene using ChIPpeakAnno (100). Motif analysis on cocaine DARs was performed using Homer (89). Gene enrichment analyses were performed using gProfiler (90) and Reactome enrichment was performed in Cytoscape (93). Track plots of ATAC-seq data were generated using SparK (94). Heatmaps and histograms of ATAC-seq reads within cFos-and ΔFosB-bound regions were generated using ngs.plot (101). Read numbers and quality control metrics for ATAC-seq libraries are shown in **Supplementary Table 7**.

### Statistical Analysis

For the early-life stress CPP experiment, we performed a three-way analysis of variance (ANOVA) with group, sex, and dose as factors. To analyze the effect of the estrous cycle within control females, we performed a two-way ANOVA with dose and estrous cycle as factors. For the XO CPP experiment, we first performed a one-way ANOVA based on genotype; then, within females, we performed a two-way ANOVA with genotype and estrous cycle as factors. For all multivariate ANOVA tests, we first evaluated whether the interaction term was significant, and if it was not significant we dropped the interaction term from the model, followed by an analysis of the main effects of the factors. For the within-female XO CPP analysis, we performed post-hoc t-tests to understand the genotype by estrous cycle interaction effect. For maternal behavior scores, we performed repeated-measures ANOVA tests with day and maternal behavior as factors, and we also analyzed the average maternal behavior scores across the 6 days using t-tests. The results of these tests were considered significant at a P < 0.05. Box plots depict the 1st–3rd quartile, with the horizontal line denoting the median and the whiskers denoting 1.5 times the inter-quartile range. The analyses were performed in R version 4.2.2 and the graphs were generated using the R ggplot2 package.

## Acknowledgements

This work was supported by the National Institute of Health under Award Number R01MH123523 (to M.K.); the NARSAD Young Investigator Grant from the Brain & Behavior Research Foundation (to M.K); and the Fordham University Interdisciplinary Research Grant (to M.K. and H.C). M.S. was supported, in part, by the National Institute of Health Award Number R01HL14530. We would further like to acknowledge the resources of the Center for Epigenomics at the Albert Einstein College of Medicine. Finally, we would like to thank Lydia Tesfa for her assistance with nuclei sorting and Benjamin Hubert for his assistance with massively parallel sequencing.

## Author Contributions

D.R., I.J., F.B., and M.K. performed experiments; D.R. and M.S. performed bioinformatics analyses; D.R. and H.C. performed statistical analyses; D.R., H.C., M.S., and M.K. interpreted the data and constructed the figures; J.M.G. contributed computational resources; D.R. and M.K. wrote the article; M.K. conceived the study and directed the project; all authors commented on and approved the final version of the paper.

I.J. is currently affiliated with Animal Welfare Division, Vetsuisse Faculty, University of Bern, Bern, Switzerland.

## Data and materials availability

RNA-seq and ATAC-seq data have been deposited at the NCBI Gene Expression Omnibus under accession number GSE229696. All other relevant data supporting the key findings of this study are available within the article and its Supplementary Information files or from the corresponding author upon request.

Correspondence and requests for materials should be addressed to M.K.

## Supplementary Information

### Other Supplementary Information for this manuscript include the following

#### Supplementary Tables

**Supplementary Table 1:** Summary statistics for the cocaine CPP test (uploaded as a separate excel file)

Suppl. Table 1A:Statistics for the Group x Sex x Dose three-way ANOVA;

Suppl. Table 1B:Statistics for the within-female Estrous Cycle x Dose two-way ANOVA.

**Supplementary Table 2:** Results of gene expression analyses in the NAc (uploaded as a separate excel file)

Suppl. Table 2A:Result table for the female control (Con) vs. maternal separation (MS) group comparison;

Suppl. Table 2B:Result table for the male Con vs. MS group comparison; Suppl. Table 2C:Result table for the Group x Sex interaction analysis; Suppl. Table 2D:Result table for Con female vs. male group comparison; Suppl. Table 2E:Result table for MS female vs. male group comparison;

Suppl. Table 2F:List of genes in each of the three gene coexpression clusters;

Suppl. Table 2G:Lists of overlapping genes altered by early-life stress in this study and two separate studies.

**Supplementary Table 3:** Gene set enrichment analysis (GSEA) Enrichment Map data for Females, Males, and Group x Sex interaction genes (uploaded as a separate excel file)

**Supplementary Table 4:** Results for differential chromatin accessibility analysis in the NAc (uploaded as a separate excel file)

Suppl. Table 4A:Result table for differentially accessible regions (DARs) in the diestrus control vs. cocaine comparison;

Suppl. Table 4B:Result table for DARs in the proestrus control vs. cocaine comparison;

Suppl. Table 4C:Result table for DARs in the male control vs. cocaine comparison;

Suppl. Table 4D:Result table for DARs in the control male vs. diestrus comparison;

Suppl. Table 4E:Result table for DARs in the control male vs. proestrus comparison;

Suppl. Table 4F:Result table for DARs in the control diestrus vs. proestrus comparison;

Suppl. Table 4G:Result table for DARs in the cocaine male vs. diestrus comparison;

Suppl. Table 4H:Result table for DARs in the cocaine male vs. proestrus comparison;

Suppl. Table 4I:Result table for DARs in the cocaine diestrus vs. proestrus comparison.

**Supplementary Table 5:** Cocaine differentially accessible regions (DARs) with AP-1 binding sites and DARs overlapping ΔFosB-bound regions (uploaded as a separate excel file)

Suppl. Table 5A:DARs that gain accessibility after cocaine in diestrus with AP-1 binding sites;

Suppl. Table 5B:DARs that gain accessibility after cocaine in proestrus with AP-1 binding sites;

Suppl. Table 5C:DARs that gain accessibility after cocaine in males with AP-1 binding sites;

Suppl. Table 5D:DARs that gain accessibility after acute cocaine in females bound by ΔFosB in females after chronic cocaine;

Suppl. Table 5E:DARs that gain accessibility after acute cocaine in males bound by ΔFosB in males after chronic cocaine.

**Supplementary Table 6:** RNA-seq data basic information for the early-life stress experiment (uploaded as a separate excel file)

**Supplementary Table 7:** ATAC-seq data basic information for the acute cocaine treatment experiment (uploaded as a separate excel file)

**Supplementary Figure 1.**
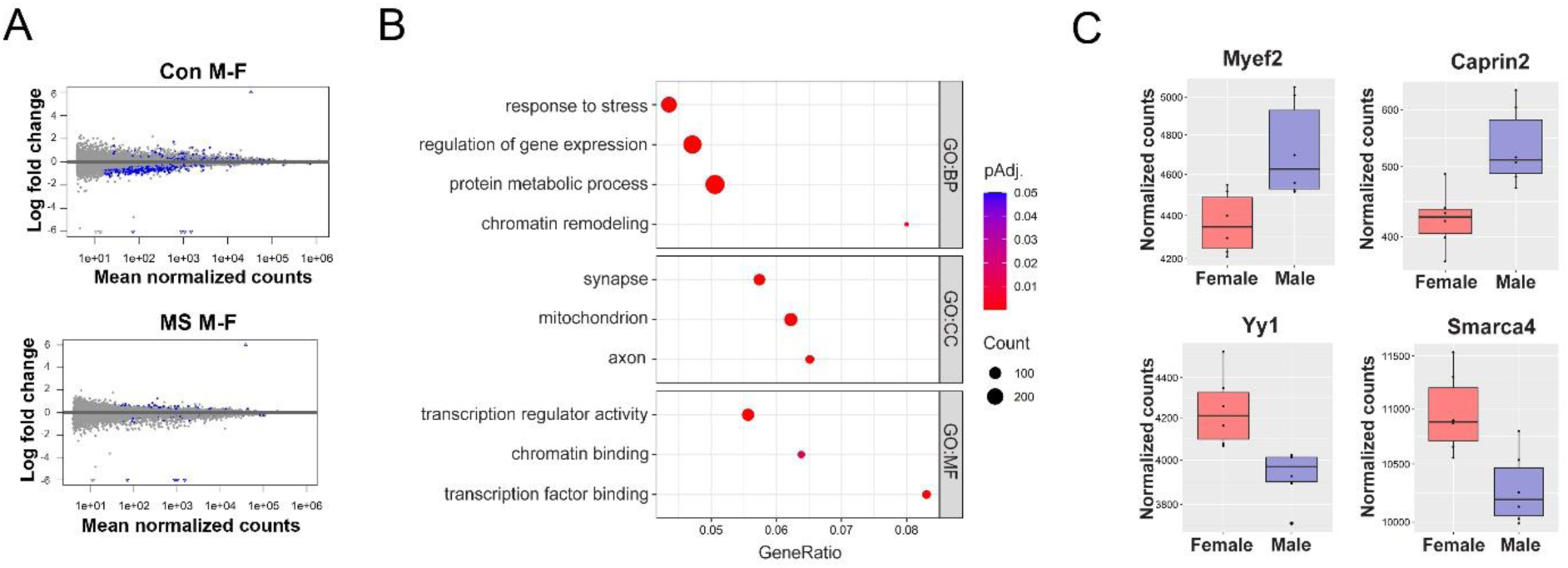
Sex differences in NAc gene expression. (**A**) RNA-seq was performed on the NAc from female (F) and male (M) mice of the control (Con) and maternal separation (MS) group (n = 6 animals/sex/group). Volcano plots depict differentially expressed genes across sex in the Con group (n = 1508), and in the MS group (n = 163); blue dots represent significant genes (p_adj_ < 0.1). (**B**) Dotplot depicting select gene ontology (GO) terms for biological process (BP), cellular compartment (CC), and molecular function (MF) significantly enriched in the genes (n=1508) differentially expressed between Con males and females in the NAc. Colors indicate adjusted *p-*values and dot size corresponds to gene count. (**C**) Example genes with differential expression across sex in the Con NAc including *Myef2* (top left) and *Caprin2* (top right), more highly expressed in males than females, as well as *Yy1* (bottom left) and *Smarca4* (bottom right), more highly expressed in females than in males. Notably, the Yy1 transcription factor is implicated in driving the transcriptional effects of early-life stress in gfemales (**Figure 2**). Box plots (box, 1^st^–3^rd^ quartile; horizontal line, median; whiskers, 1.5× IQR).

**Supplementary Figure 2.**
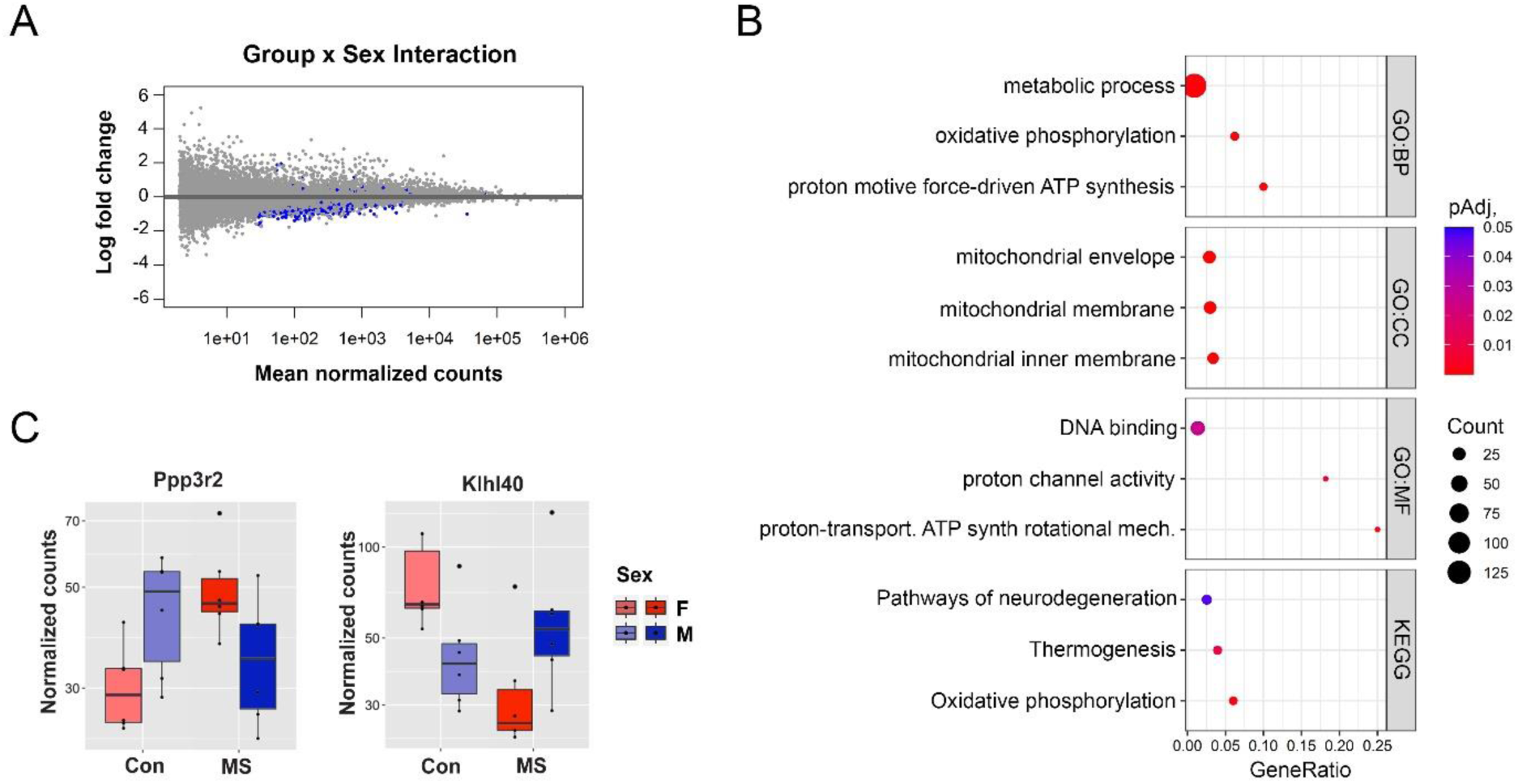
Early-life stress and sex interact to alter NAc gene expression. **(A)** Volcano plot depicts differentially expressed genes identified in the Group x Sex interaction analysis (n = 455); blue dots represent significant genes (p_adj_ < 0.1). (**B**) Dotplot depicting select gene ontology (GO) terms for biological process (BP), cellular compartment (CC), and molecular function (MF), as well as KEGG pathways, significantly enriched in genes with a significant group by sex interaction. Colors indicate adjusted *p-*values and dot size corresponds to gene count. (**C**) Example genes include *Ppp3r2* (left), which is upregulated in females but downregulated in males after early-life stress, and *Klhl40* (right), which is downregulated in females but upregulated in males after early-life stress. Box plots (box, 1^st^–3^rd^ quartile; horizontal line, median; whiskers, 1.5× IQR). Con, control group; MS, maternal separation group.

**Supplementary Figure 3.**
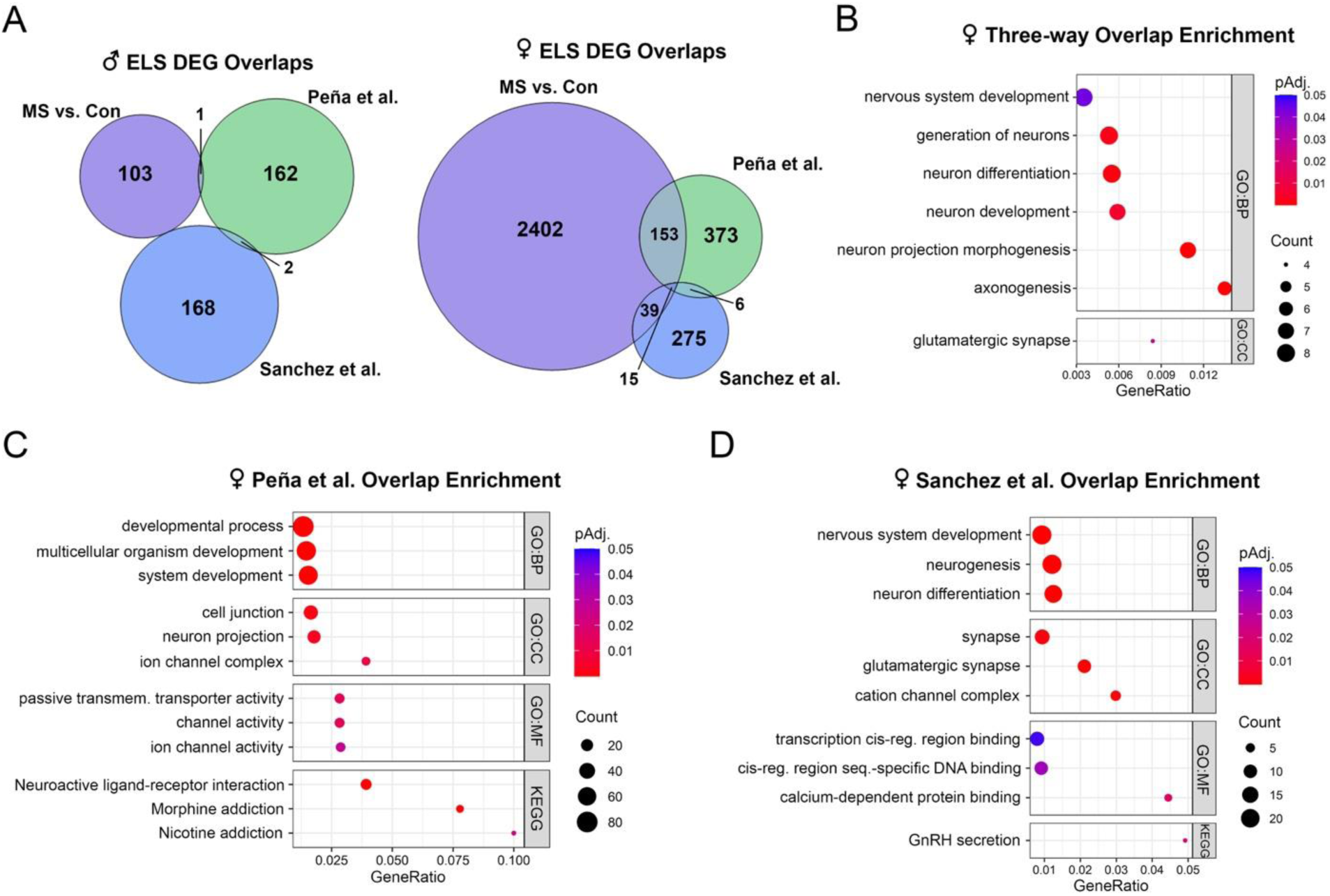
Shared effects of early-life stress on female NAc gene expression across study paradigms. (**A**) Venn diagrams showing the overlap of differentially expressed genes (DEGs) altered by early-life stress in the NAc in this study (MS vs. Con), a study utilizing limited-bedding and maternal separation from postnatal day (P) 10-17 (Peña et al.), and a study utilizing limited nesting and bedding from P2-P10 (Sanchez et al.) in males (left) and females (right). (**B**) Dotplot depicting select gene ontology (GO) terms for biological process (BP) and cellular compartment (CC) significantly enriched in female DEGs shared between the three studies. (**C**) Dotplot depicting select GO terms for BP, CC, and molecular function (MF), as well as KEGG pathways, significantly enriched in female DEGs shared between this study and the Peña et al. study. (**D**) Dotplot depicting select GO terms for BP, CC, and MF, as well as KEGG pathways, significantly enriched in female DEGs shared between this study and the Sanchez et al. study. Dotplot colors indicate adjusted *p-*values and dot size corresponds to gene count. Con, control group; MS, maternal separation group.

**Supplementary Figure 4.**
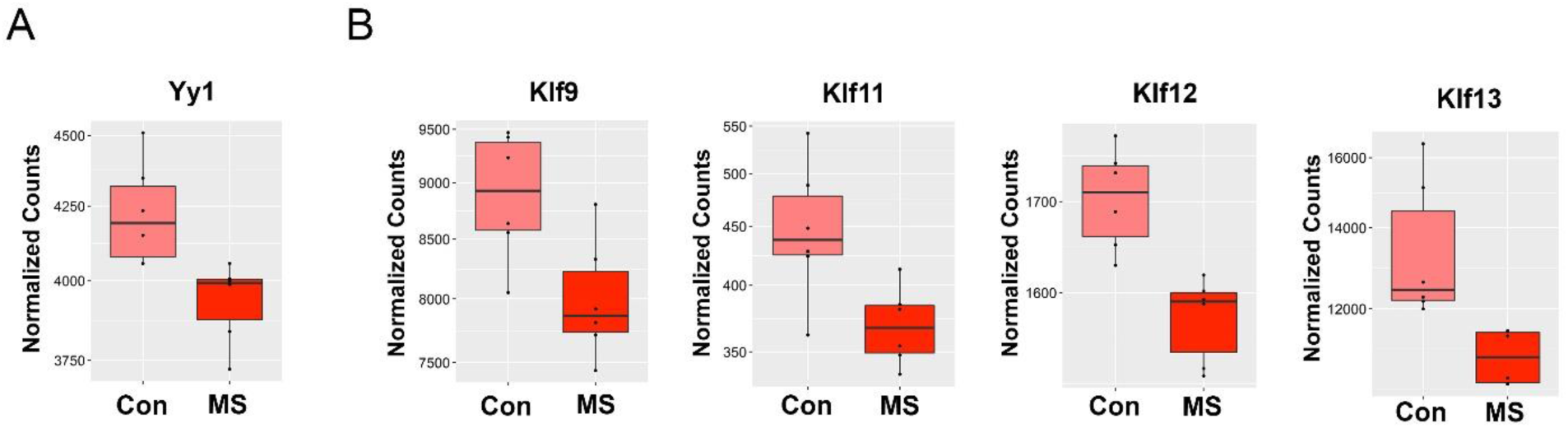
Yy1 and Klf-family transcription factors are downregulated in the female NAc by early-life stress. (**A**) Normalized count plot of the gene encoding the Yy1 transcription factor, whose binding sites are enriched in promoters of cluster 1 genes (**Figure 2**) are shown for the female group. (**B**) Normalized count plots of genes encoding Klf-family transcription factors, including *Klf9*, *Klf11*, *Klf12*, and *Klf13*, whose binding sites are enriched in promoters of cluster 3 genes (**Figure 2**) are shown for the female group. Box plots (box, 1^st^–3^rd^ quartile; horizontal line, median; whiskers, 1.5× IQR). Con, control group; MS, maternal separation group.

**Supplementary Figure 5.**
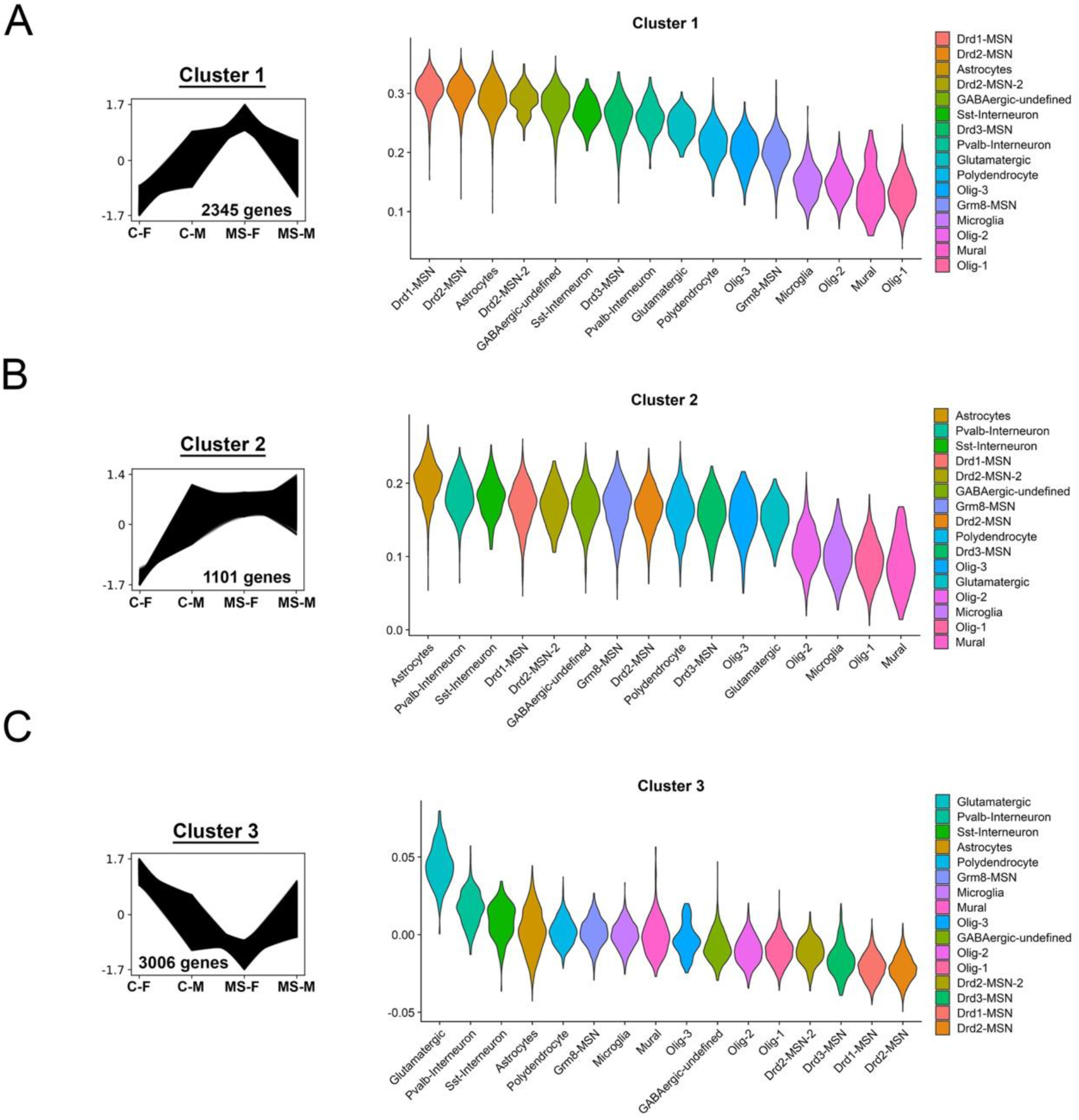
Cell type-biased expression patterns of coexpression cluster genes. (**A**) Genes in cluster 1 (left, **Figure 2**) are primarily expressed in dopamine receptor 1-(Drd1), and 2-(Drd2), expressing medium spiny neurons (MSNs), as well as astrocytes, as shown in the violin plots (right). (**B**) Genes in cluster 2 (left, **Figure 2**) are primarily expressed in astrocytes and parvalbumin-(Pvalb) and somatostatin-(Sst) expressing interneurons (MSNs), as shown in the violin plots (right). (**C**) Genes in cluster 3 (left, **Figure 2**) are primarily expressed in glutamatergic neurons and parvalbumin-(Pvalb) and somatostatin-(Sst) expressing interneurons (MSNs), as shown in the violin plots (right). C-F, control female; C-M, control male; MS-F, maternal separation female; MS-M, maternal separation male.

**Supplementary Figure 6.**
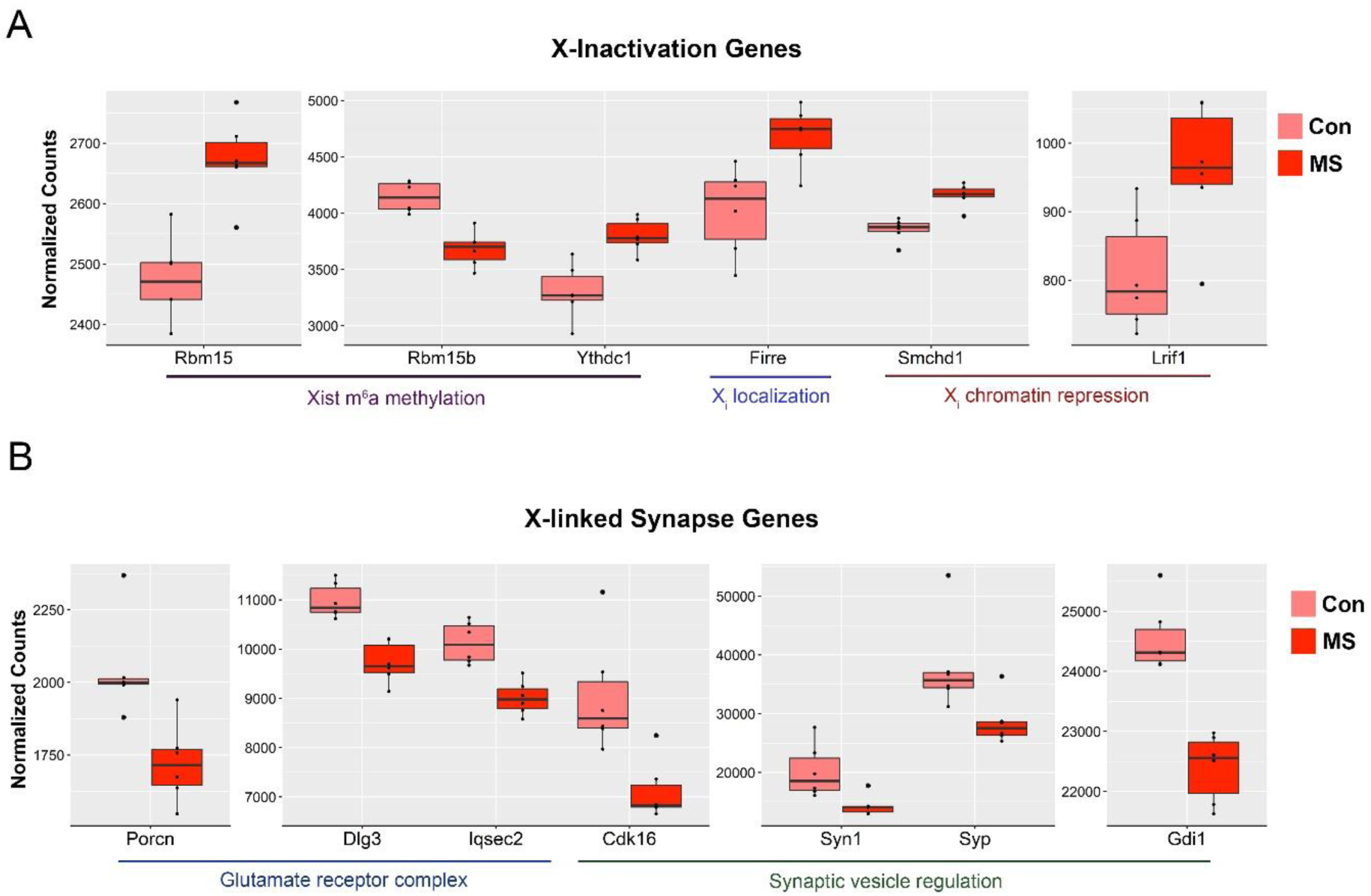
Early life stress alters genes involved in X-inactivation and X-linked genes involved in synaptic function in the female Nac. (**A**) Normalized count plots showing the altered expression of autosomal and X-linked genes involved in X-inactivation, including Xist m^6^a methylation (*Rbm15*, *Rbm15b*, and *Ythdc1*; autosomal), localization of the inactive-X chromosome (X_i_) (*Firre*; X-linked), and X_i_ chromatin repression (*Smchd1* and *Lrif1*; autosomal; left to right) (**B**) Normalized count plots showing the altered expression of X-linked genes involved in synapse function, including genes encoding members of the glutamate receptor complex (*Porcn*, *Dlg3*, and *Iqsec2*) and genes involved in synaptic vesicle regulation (*Cdk16*, *Syn1*, *Syp*, and *Gdi1*; left to right). Box plots (box, 1st–3rd quartile; horizontal line, median; whiskers, 1.5× IQR). Con, control group; MS, maternal separation group.

**Supplementary Figure 7.**
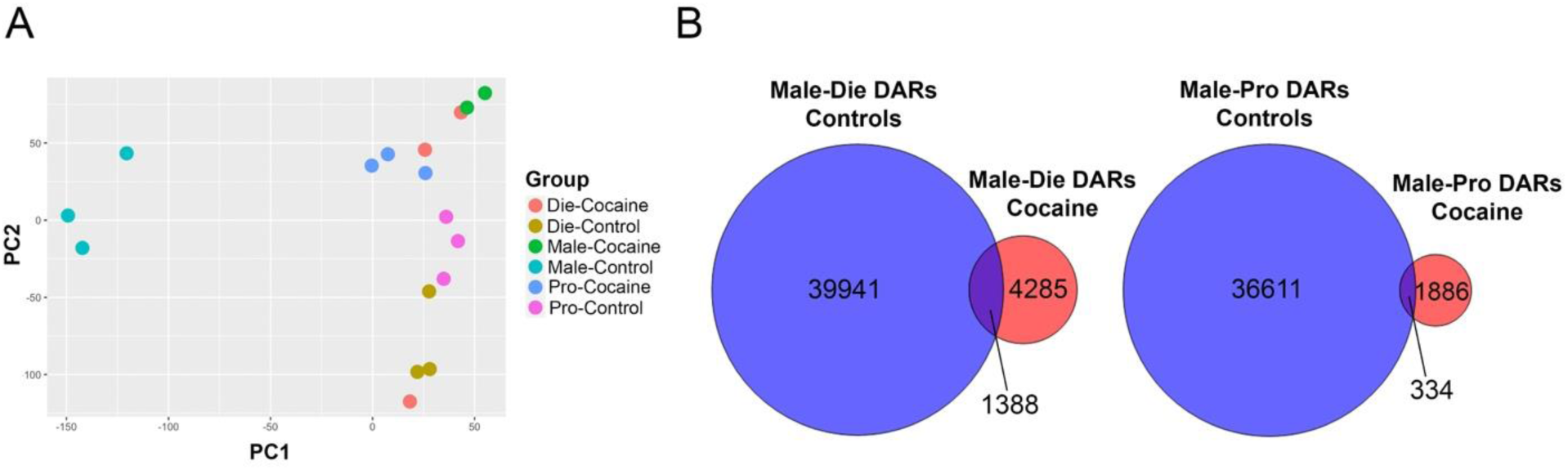
Acute cocaine exposure reduces sex differences in NAc chromatin accessibility. (**A**) A PCA plot showing the clustering of ATAC-seq replicates across groups and treatment. While male and female controls are separated along the first principal component (PC1), cocaine-treated males and females cluster together more closely. (**B**) Venn diagrams depicting the overlap of differentially accessible chromatin regions (DARs) between males and diestrus females in the control and cocaine conditions (left) and between males and proestrus females in the control and cocaine conditions (right), illustrating that cocaine treatment produces a smaller, largely distinct set of sex DARs compared to those present in controls. Die, diestrus; Pro, proestrus; Male, males.

**Supplementary Figure 8.**
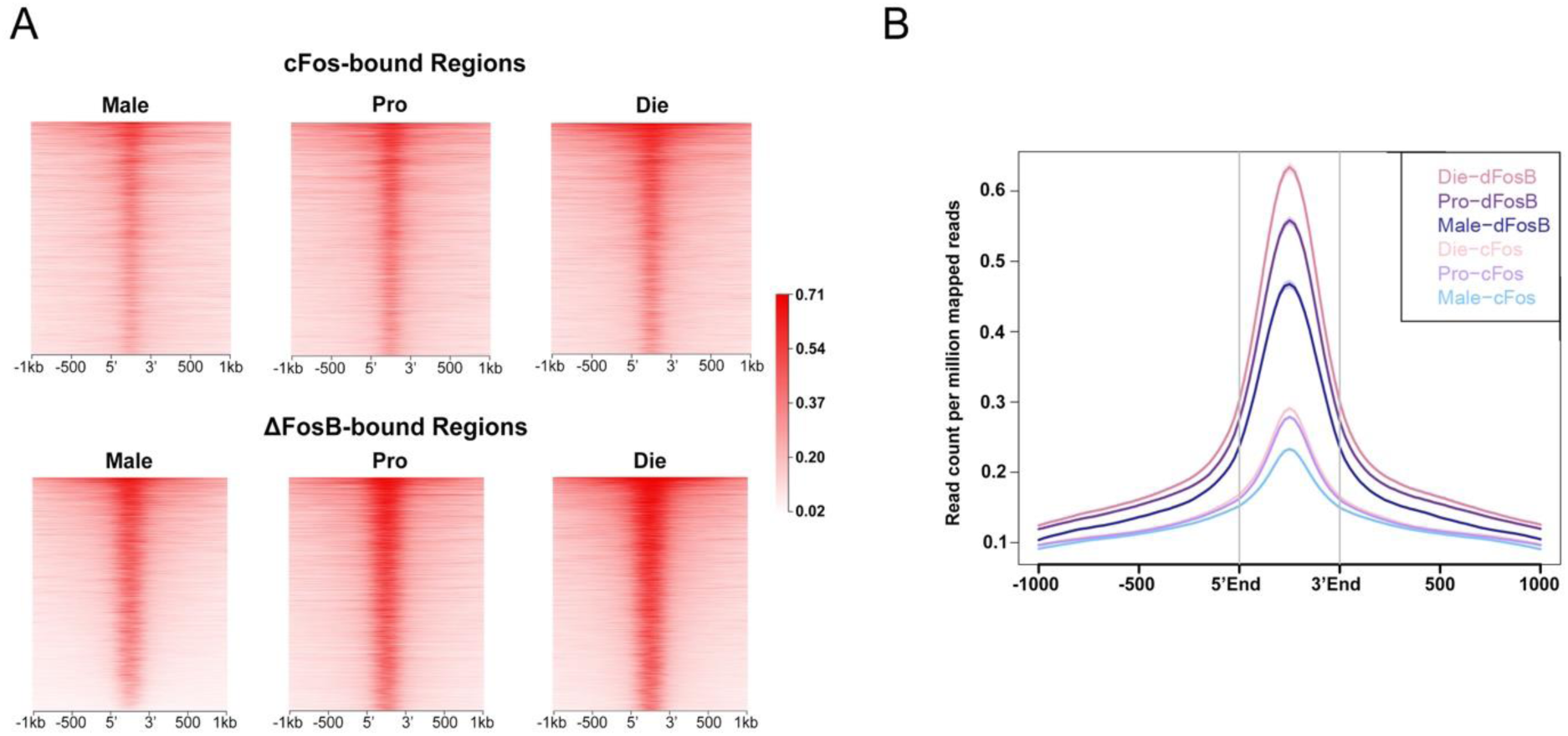
Accessible chromatin after acute cocaine in the NAc overlaps more strongly with ΔFosB-bound regions than cFos-bound regions. (**A**) Heatmaps show the density of ATAC-seq reads in acute cocaine treated males (left) proestrus females (middle), and diestrus females (right) within regions that are bound by cFos (top) or ΔFosB (bottom). (**B**) A histogram representation of the data shown in (**A**), demonstrating a higher ATAC-seq signal in ΔFosB-bound regions than cFos-bound regions for all groups, especially diestrus females. Die, diestrus; Pro, proestrus; Male, males.

**Supplementary Figure 9.**
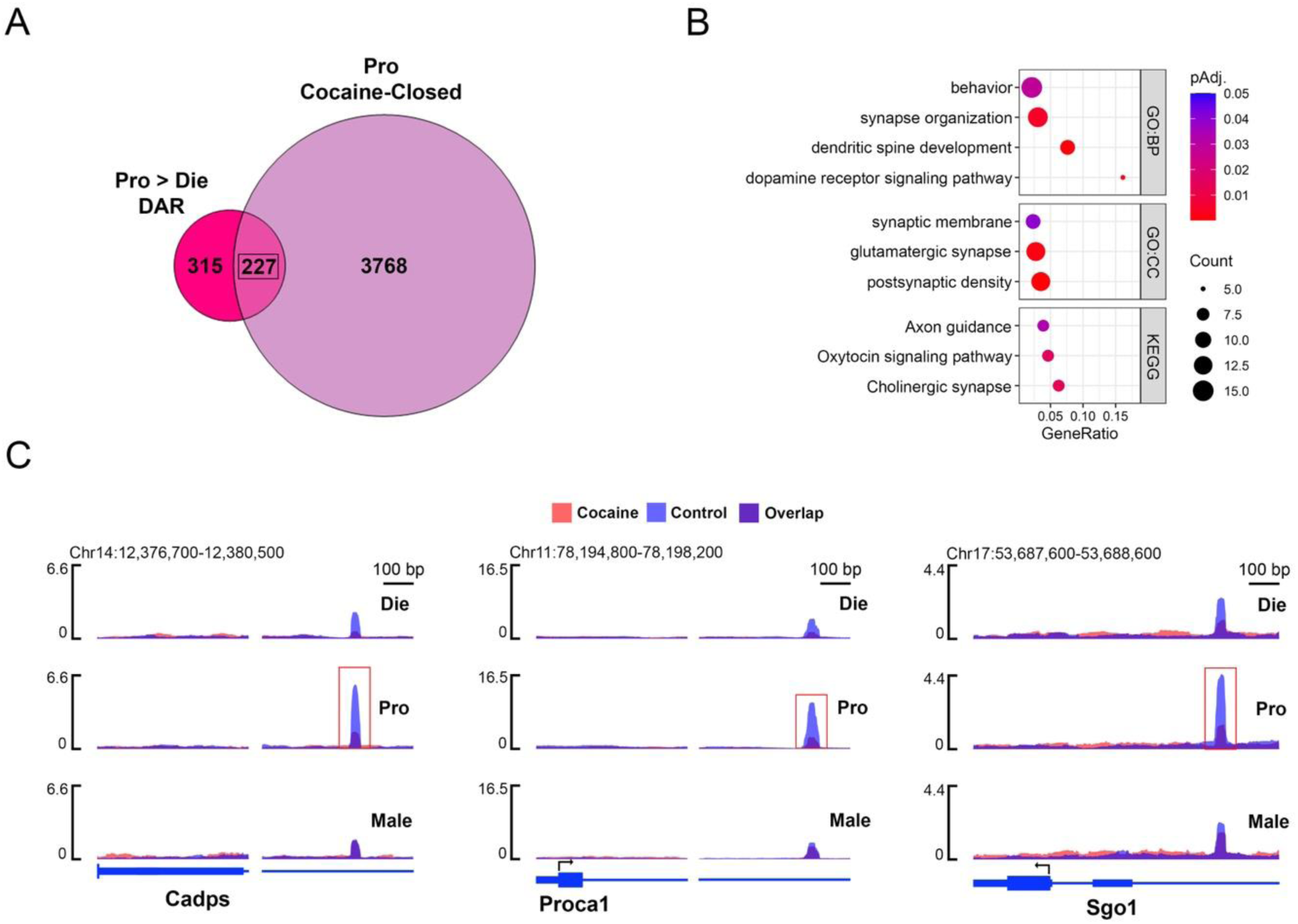
Regions more accessible in proestrus female controls become less accessible after acute cocaine exposure. (**A**) A Venn diagram showing the overlap between differentially accessible regions (DARs) that are more accessible in proestrus than diestrus controls, with regions that are less accessible in proestrus females following acute cocaine exposure. (**B**) A dotplot depicting select gene ontology (GO) terms for biological process (BP) and cellular component (CC), as well as KEGG pathways, significantly enriched in the genes annotated to overlapping regions highlighted in (**A**). Dotplot colors indicate adjusted *p-*values and dot size corresponds to gene count. (**C**) SparK plots of group-average normalized ATAC-seq reads (n = 2-3 replicates or 6-9 animals/group) are shown for example regions from the overlap shown in (**A**), including regions near the transcription start sites (TSSs) of *Cadps* (left), *Proca1* (middle), and *Sgo1* (right), all of which are more accessible in proestrus controls before cocaine but lose accessibility in this group after cocaine. Die, diestrus; Pro, proestrus; Male, males.

**Supplementary Figure 10.**
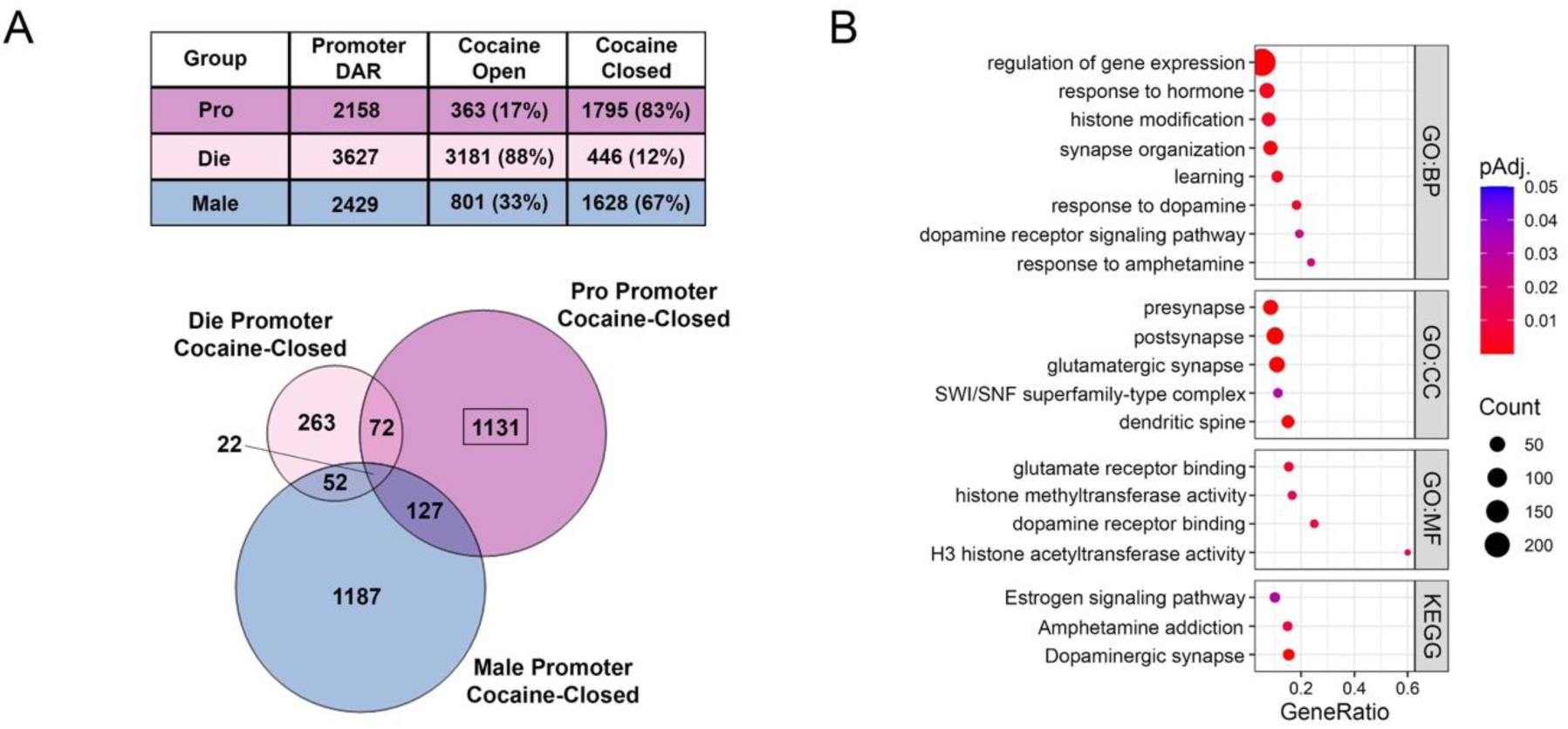
Chromatin surrounding gene promoters are less accessible after acute cocaine in proestrus females. (**A**) A table showing the number of cocaine differentially accessible regions (DARs) that overlap gene promoters and the number of these regions that become more (Cocaine Open) or less (Cocaine Closed) accessible after cocaine in each group (top), as well as a Venn diagram showing the overlap of promoter DARs that are less accessible after cocaine in all three groups (bottom). (**B**) A dotplot depicting select gene ontology (GO) terms for biological process (BP), cellular component (CC), and molecular function (MF) as well as KEGG pathways, significantly enriched in the genes annotated to proestrus-specific promoter DARs that are less accessible after cocaine, highlighted in the Venn diagram shown in (**A**). Dotplot colors indicate adjusted *p-*values and dot size corresponds to gene count.

**Supplementary Figure 11.**
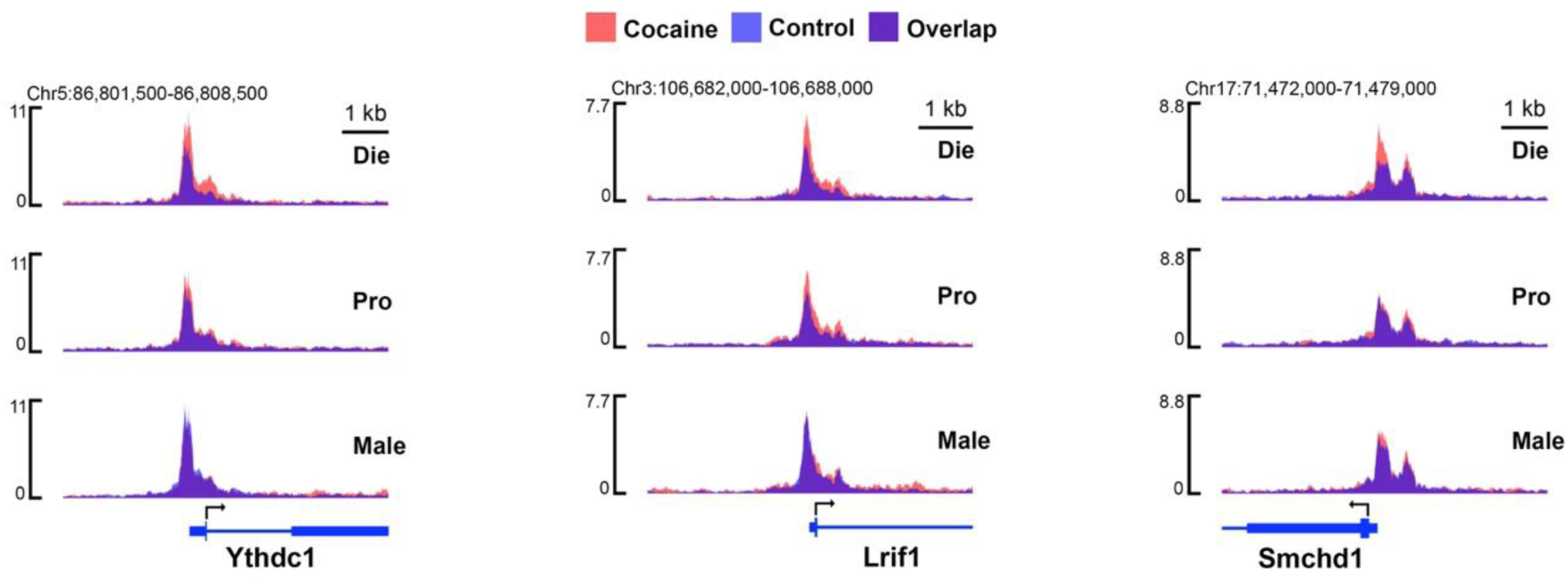
Promoter regions of autosomal genes involved in X-inactivation are more accessible after cocaine in diestrus females. SparK plots of group-average normalized ATAC-seq reads (n = 2-3 replicates or 6-9 animals/group) are shown for autosomal genes involved in X-inactivation, whose expression is also altered in females by early-life stress (**Suppl. Figure 6A**), including regions overlapping the transcription start sites (TSSs) of *Ythdc1* (left), *Lrif1* (middle), and *Smchd1* (right), all of which are more accessible after acute cocaine specifically in diestrus females. Die, diestrus; Pro, proestrus; Male, males.

**Supplementary Figure 12.**
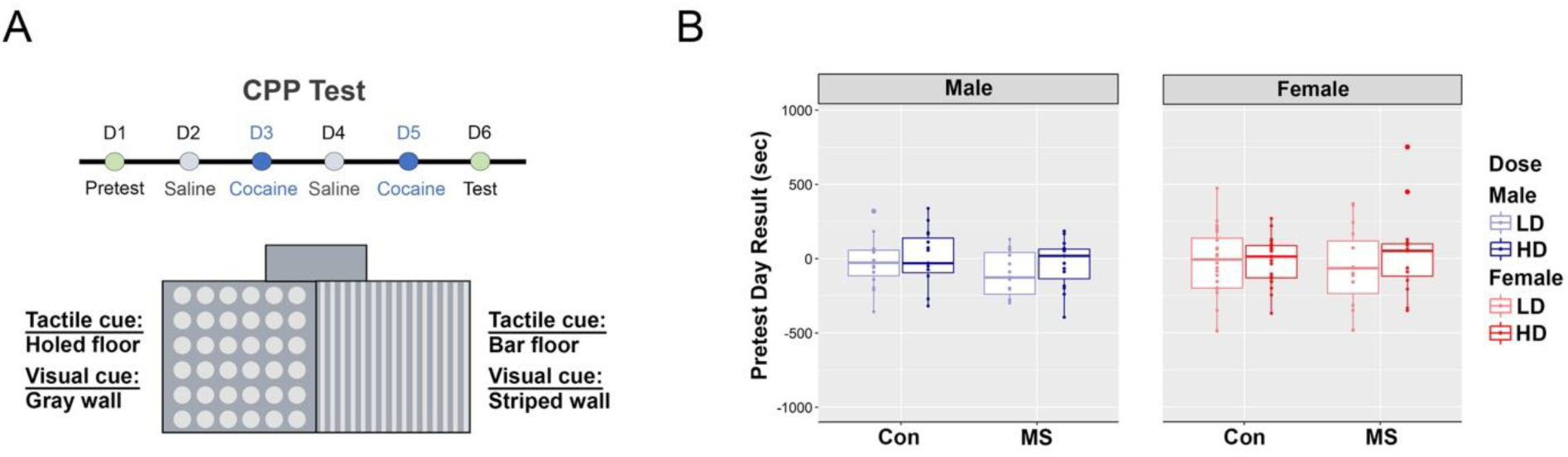
Cocaine conditioned place preference (CPP) paradigm. (**A**) A schematic showing the timeline of the cocaine CPP test, which includes a pretest day, four conditioning days alternating between saline and cocaine treatment, and a test day (top), as well as an illustration of the CPP apparatus which has two chambers separated by a closeable corridor which are distinguishable by visual and tactile cues (bottom). (**B**) Plots showing the time spent in each compartment of the CPP apparatus by all animals across sex, group, and dose during the pretest day, demonstrating that animals in the experiment had no overall preference for either compartment. Box plots (box, 1st–3rd quartile; horizontal line, median; whiskers, 1.5× IQR). D, Day; Con, control group; MS, maternal separation group; LD, low-dose; HD, high-dose.

**Supplementary Figure 13.**
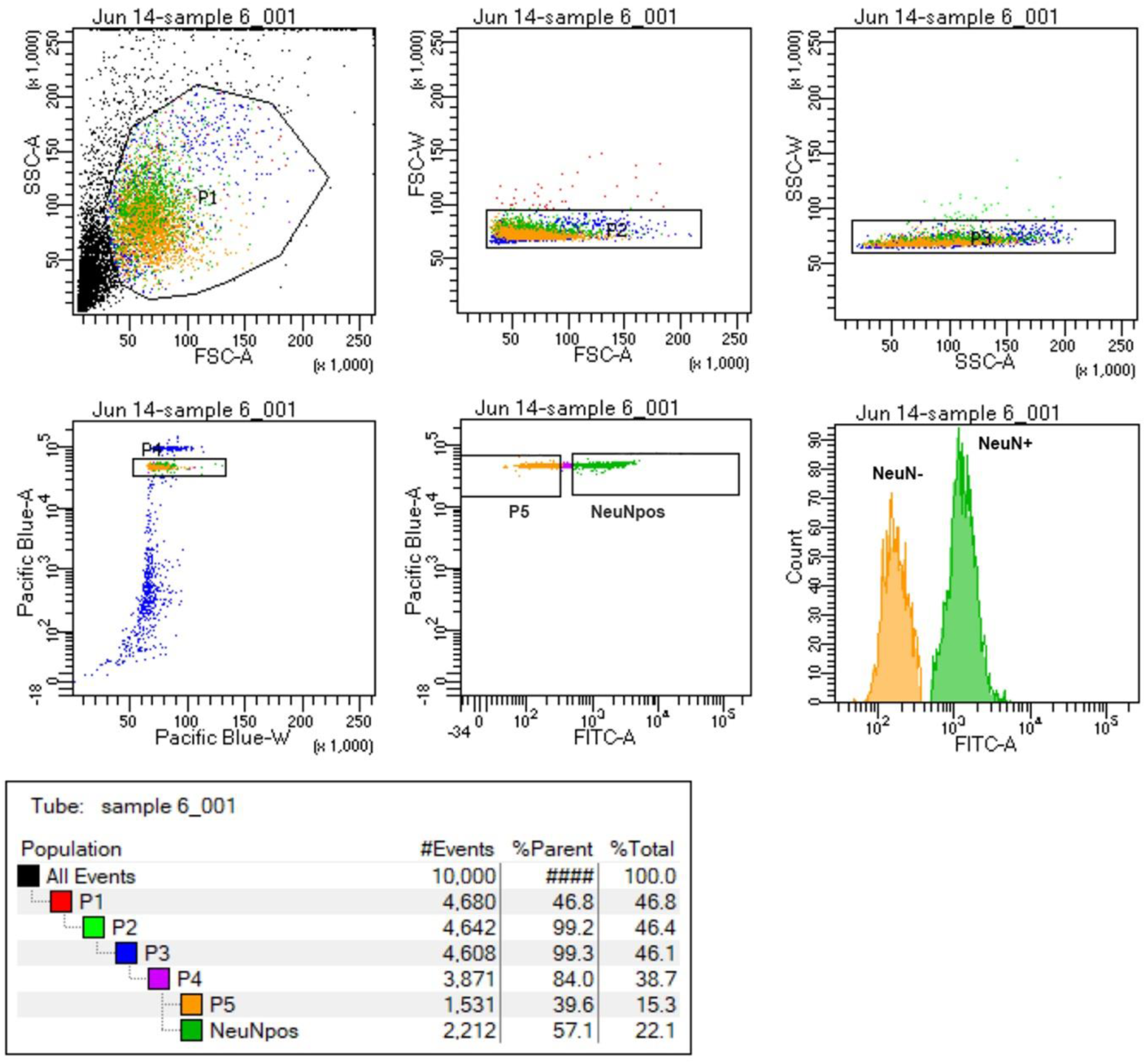
Purification of neuronal (NeuN+) nuclei with fluorescence-activated nuclei sorting (FANS). A representative FANS report demonstrating the gating procedure that allowed:1) the separation of nuclei from cellular debris (P1-P3); 2) the separation of single, intact nuclei using the DAPI signal (P4); and 3) a specific purification of neuronal nuclei with a NeuN+ signal (P6) from non-neuronal (NeuN-) nuclei (P5).

